# Functional proteomic atlas of HIV-infection in primary human CD4+ T cells

**DOI:** 10.1101/389346

**Authors:** Adi Naamati, James C Williamson, Edward JD Greenwood, Sara Marelli, Paul J Lehner, Nicholas J Matheson

## Abstract

Viruses manipulate host cells to enhance their replication, and the identification of host factors targeted by viruses has led to key insights in both viral pathogenesis and cellular physiology. We previously described global changes in cellular protein levels during human immunodeficiency virus (HIV) infection using transformed CEM-T4 T cells as a model. In this study, we develop an HIV reporter virus displaying a streptavidin-binding affinity tag at the surface of infected cells, allowing facile one-step selection with streptavidin-conjugated magnetic beads. We use this system to obtain pure populations of HIV-infected primary human CD4+ T cells for detailed proteomic analysis, including quantitation of >9,000 proteins across 4 different donors, and temporal profiling during T cell activation. Remarkably, amongst 650 cellular proteins significantly perturbed during HIV infection of primary T cells (q<0.05), almost 50% are regulated directly or indirectly by the viral accessory proteins Vpr, Vif, Nef and Vpu. The remainder have not been previously characterised, but include novel Vif-dependent targets FMR1 and DPH7, and 192 targets not identified and/or regulated in T cell lines, such as AIRD5A and PTPN22. We therefore provide a high-coverage functional proteomic atlas of HIV infection, and a mechanistic account of HIV-dependent changes in its natural target cell.

---

Remodelling of the host proteome during viral infection may reflect direct effects of viral proteins, secondary effects or cytopathicity accompanying viral replication, or host countermeasures such as the interferon (IFN) response. By defining time-dependent changes in protein levels in infected cells, and correlating temporal profiles of cellular and viral proteins, we have shown that it is possible to differentiate these phenomena, and identify direct cellular targets of human cytomegalovirus (HCMV) and HIV^1–3^. To enable time course analysis and minimise confounding effects from uninfected bystander cells, pure populations of synchronously infected cells must be sampled sequentially as they progress through a single round of viral replication. In the case of HIV, we previously satisfied these conditions by spinoculating the highly permissive CEM-T4 lymphoblastoid T cell line^4–6^ with Env-deficient NL4-3-ΔEnv-EGFP virus^7^ at a high multiplicity of infection (MOI)^3^.

The utility of cancer cell line models (such as CEM-T4) is, however, limited by the extent to which they retain the characteristics of the primary cells from which they were derived, and cancer-specific and *in vitro* culture-dependent reprogramming are well described^8^. For example, the HIV accessory proteins Vif, Nef and Vpu are required for viral replication in primary T cells, but not in many T cell lines^9–12^, and HIV is restricted by type I interferon (IFN) in primary T cells, but not CEM-derived T cells^13^. In addition, whilst ensuring a high % infection, dysregulation of the cellular proteome at high MOIs may not be indicative of protein changes when a single transcriptionally active provirus is present per cell. In this study, we therefore sought to apply our temporal proteomic approach to HIV infection of primary human CD4+ T lymphocytes, the principle cell type infected *in vivo*, at an MOI <1. To this end, we have developed a reporter virus for antibody-free magnetic cell sorting (AFMACS)^14^ of HIV-infected cells (**Fig. 1a**). AFMACS allows facile, one-step affinity purification of infected cells from mixed cultures, bypassing the need for high MOIs or fluorescence-activated cell sorting (FACS). We use this system to generate a detailed atlas of cellular protein dynamics in HIV-infected primary human CD4+ T cells, show how this resource can be used to identify novel cellular proteins regulated by HIV, and assign causality to individual HIV accessory proteins.

**Figure 1:**
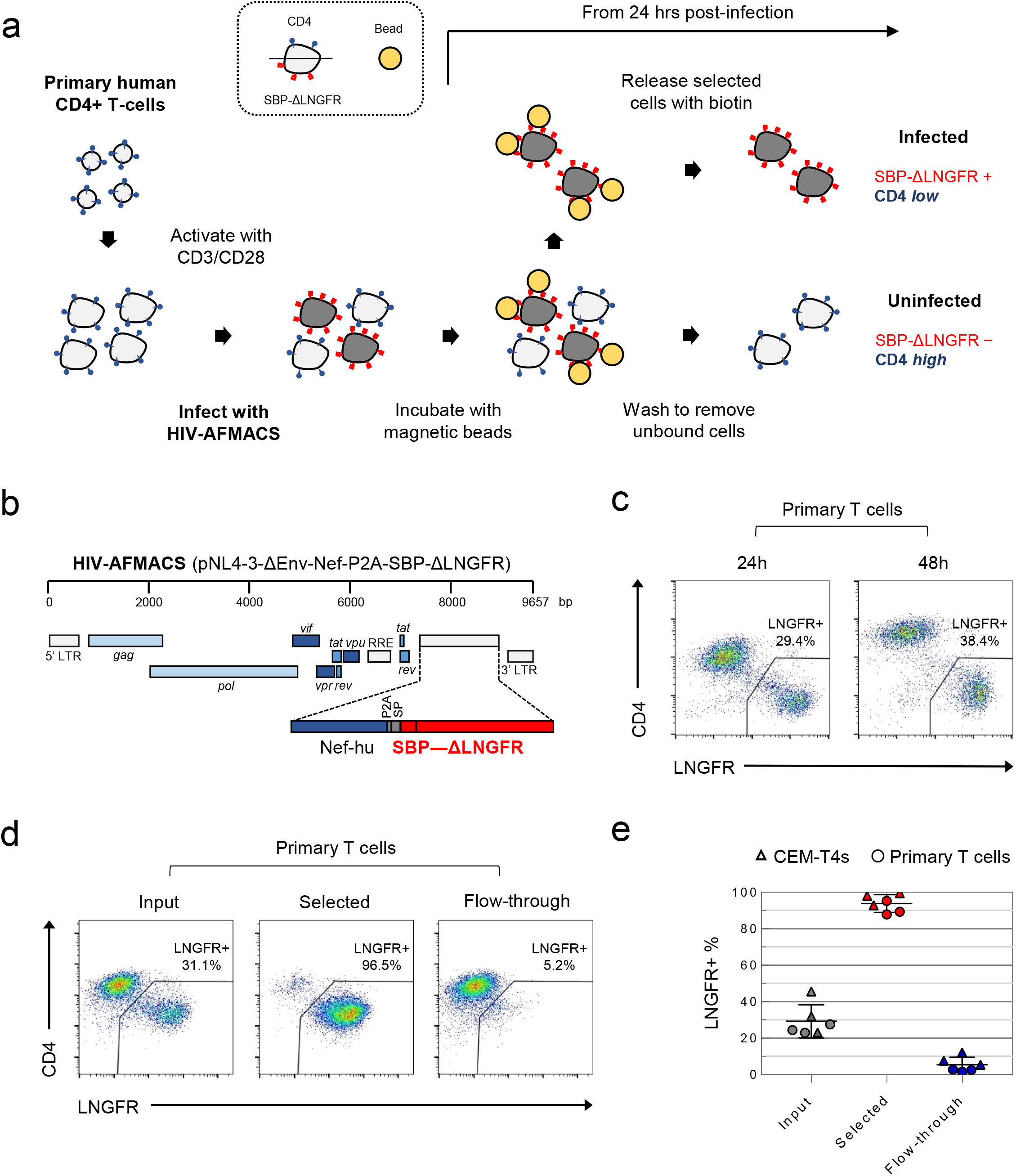
Antibody-free magnetic selection of HIV-infected primary T cells. **a**, Workflow for AFMACS-based magnetic selection of HIV-infected primary T cells. **b**, Schematic of HIV-AFMACS provirus (pNL4-3-ΔEnv-Nef-P2A-SBP-ΔLNGFR). For simplicity, reading frames are drawn to match the HXB2 HIV-1 reference genome. Length is indicated in base pairs (bp). The complete sequence is available in the **Supplementary Methods**. Nef-hu, codon-optimised Nef; RRE, Rev response element; SP, signal peptide. **c**, Expression of cell surface SBP-ΔLNGFR and CD4 on primary T cells 24 or 48 hrs post-infection with HIV-AFMACS. Cells were stained with anti-LNGFR and anti-CD4 antibodies at the indicated time points and analysed by flow cytometry. **d**, **e**, Magnetic sorting of HIV-infected (red, LNGFR+, CD4 low) and uninfected (blue, LNGFR-, CD4 high) cells. Cells were separated using AFMACS 48 hrs post-infection with HIV-AFMACS and analysed as in c. Representative (**d**) and summary (**e**) data from 6 independent experiments in CEM-T4s (triangles) and primary T cells (circles) are shown, with means and 95% confidence intervals (CIs).

AFMACS-based magnetic selection requires the high-affinity 38 amino acid streptavidin-binding peptide (SBP)^15^ to be displayed at the cell surface by fusion to the N-terminus of the truncated Low-affinity Nerve Growth Factor Receptor (SBP-ΔLNGFR)^16^. Cells expressing this marker may be selected directly with streptavidin-conjugated magnetic beads, washed to remove unbound cells, then released by incubation with the naturally occurring vitamin biotin^14^. To engineer a single round HIV reporter virus encoding SBP-ΔLNGFR, we considered 3 settings in the proviral construct: fused to the endogenous Env signal peptide (as a direct replacement for Env); or as an additional cistron, downstream of *nef* and either a P2A peptide or IRES. We used Env-deficient pNL4-3-ΔEnv-EGFP (HIV-1) as a backbone and, since increased size of lentiviral genome is known to reduce packaging efficiency^17^, tested each approach in constructs from which EGFP was removed and/or the 3’ long terminal repeat (LTR) truncated. Further details relating to construct design are described in the **Methods** and **Supplementary Methods**.

For initial screening, VSVg-pseudotyped viruses were made in 293T cells under standard conditions, and used to spinoculate CEM-T4 T cells (CEM-T4s). Infected cells were identified by expression of EGFP and/or cell surface LNGFR, combined with Nef/Vpu-mediated downregulation of CD4^18,19^. Whilst infection is not truly “productive” (because Env is deleted), Gag alone is sufficient for assembly and release of virions^20^, and other structural and non-structural viral proteins are expressed in accordance with full length viral infection^3^. As expected, all viruses tested expressed SBP-ΔLNGFR at the cell surface of infected cells (**Supplementary Fig. 1a**), but the larger constructs resulted in lower infectious viral titres (**Supplementary Fig. 1a,b**). We therefore selected pNL4-3-ΔEnv-SBP-ΔLNGFR, pNL4-3-ΔENV-Nef-P2A-SBP-ΔLNGFR and pNL4-3-ΔEnv-Nef-IRES-SBP-ΔLNGFR-Δ3 for further evaluation (**Supplementary Fig. 2a**). Viruses generated from these constructs expressed high levels of SBP-ΔLNGFR 48 hrs post-infection, and depleted CD4 and tetherin to a similar extent. However, only the pNL4-3-ΔENV-Nef-P2A-SBP-ΔLNGFR virus (**Fig. 1b**) expressed high levels of LNGFR 24 hrs post-infection in both CEM-T4s (**Supplementary Fig. 2a**) and primary human CD4+ T cells (**Fig. 1c**). This is consistent with Nef-P2A-SBP-ΔLNGFR expression from completely spliced transcripts early in HIV infection^21^, with the P2A peptide ensuring that translation of Nef and SBP-ΔLNGFR follow similar kinetics.

Since analysis of cells at early as well as late time points is essential for the generation of time course data, we focussed on pNL4-3-ΔEnv-Nef-P2A-SBP-ΔLNGFR (now termed HIV-AFMACS). To confirm that HIV-AFMACS virus could be used for cell selection (**Fig. 1a**), infected primary T cells were captured by streptavidin-conjugated magnetic beads, released by incubation with excess biotin, then analysed by flow cytometry. Compared with unselected cells (input) or cells released during washing (flow-through), selected cells were markedly enriched for SBP-ΔLNGFR expression and CD4 downregulation (**Fig. 1d**). In fact, from mixed populations containing approximately 20-40% infected cells (corresponding to an effective MOI</=0.5), purities of >/=90% were routinely achieved by AFMACS of both CEM-T4s and primary human CD4+ T cells, with </=10% infected cells lost in the flow-through (**Fig. 1e**).

To gain a comprehensive, unbiased overview of viral and cellular protein dynamics during HIV-infection of its natural target cell, we used the HIV-AFMACS virus to spinoculate activated, primary human CD4+ T cells, sorted infected (SBP-ΔLNGFR positive) and uninfected (SBP-ΔLNGFR negative) cells by AFMACS 24 hrs and 48 hrs post-infection, and analysed whole cell lysates using tandem mass tag (TMT)-based quantitative proteomics (**Fig. 2a,b and Supplementary Fig. S3a**)^1,3^. Interpretation of HIV-dependent proteomic remodelling in primary T cells is complicated by concurrent changes in relative protein abundance resulting from T cell activation^22^. We therefore exploited multiplex TMT labelling to measure parallel protein abundances in resting and activated (uninfected) T cells from the same donor, as well as control (mock-infected) T cells obtained at each time point.

**Figure 2:**
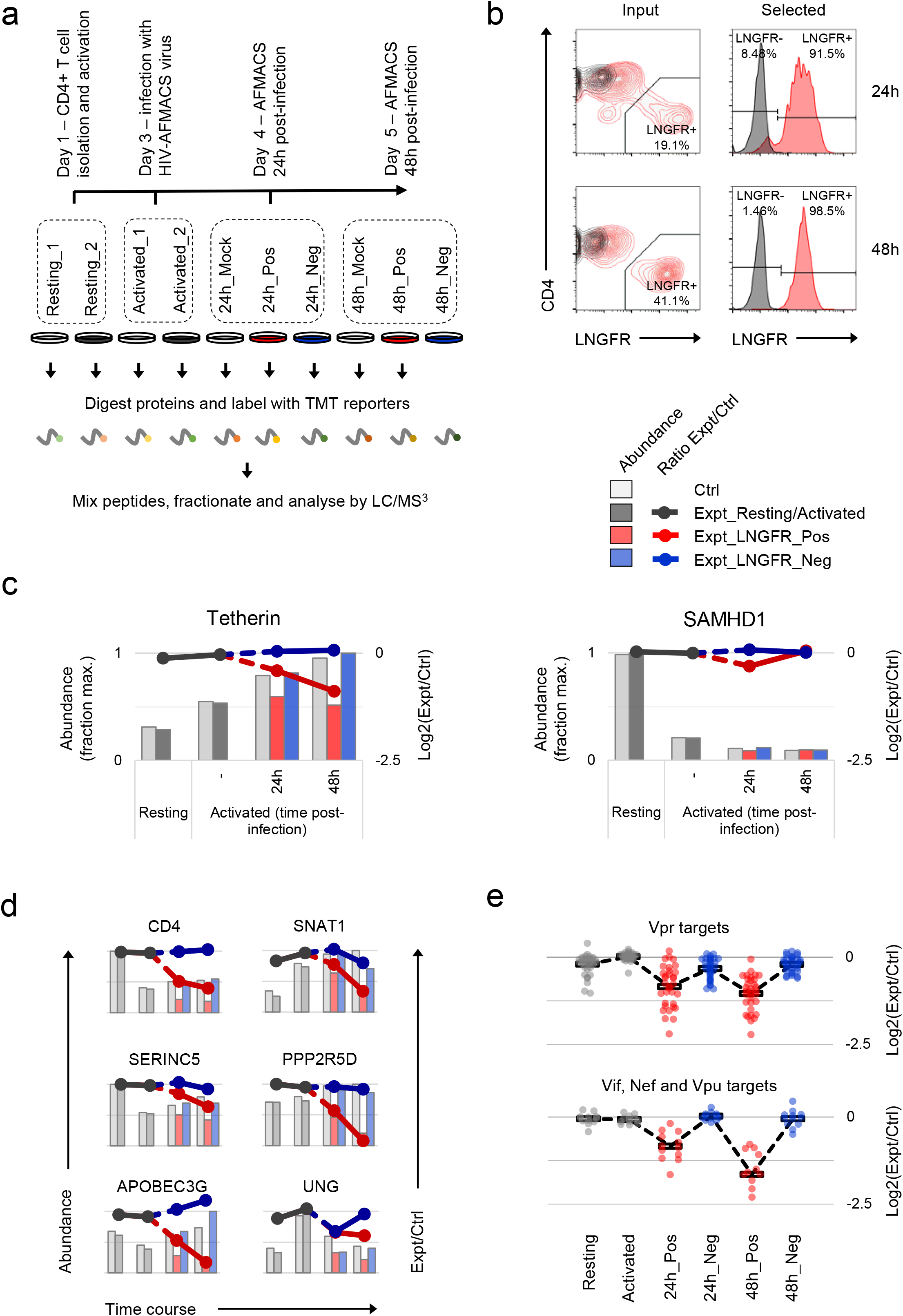
Temporal proteomic analysis of HIV infection in primary T cells. **a**, Overview of time course proteomic experiment. Control (pale grey, resting/activated/mock) and experimental (dark grey, resting/activated; red, LNGFR+, HIV-infected; blue, LNGFR-, uninfected) cells are indicated for each condition/time point. **b**, Magnetic sorting of HIV-infected (LNGFR+, selected) cells used for **a**. Corresponding uninfected (LNGFR-, flow-through) cells are shown in **Supplementary Fig. 3a**. Cells were separated using AFMACS at the indicated time points post-infection with HIV-AFMACS, stained with anti-LNGFR and anti-CD4 antibodies and analysed by flow cytometry. **c**, Expression profiles of illustrative restriction factors regulated by T cell activation and HIV infection (tetherin) or T cell activation alone (SAMHD1) in cells from **a,b**. Relative abundances (bars, fraction of maximum) and log2(ratio)s of abundances (lines) in experimental (Expt):control (Ctrl) cells are shown for each condition/time point and coloured as in **a** (summarised in the legend). **d**, Expression profiles of illustrative accessory protein targets (CD4, Nef/Vpu; SERINC5, Nef; SNAT1, Vpu; APOBEC3G, Vif; PPP2R5D, Vif; UNG, Vpr) in cells from **a,b**. Axes, scales and colours are as in **c**. Expression profiles of other accessory protein targets are shown in **Supplementary Fig. 3b**. **e**, Patterns of temporal regulation of Vpr vs other accessory protein (Vif/Nef/Vpu) targets in cells from **a,b**. Log2(ratio)s of abundances in experimental (Expt):control (Ctrl) cells are shown for 45 accessory protein targets (as in **Supplementary Fig. 7b**). Colours are as in c, and average profiles (mean, black lozenges/dotted lines) are highlighted for each group of targets.

In total, we quantitated 9,076 proteins across 10 different conditions. This and other datasets described in this study will be deposited to the ProteomeXchange consortium (accessible at http://proteomecentral.proteomexchange.org) and are summarised in an interactive spreadsheet which allows generation of temporal profiles for any quantitated proteins of interest (**Supplementary Table 1**). For example, the restriction factor tetherin (targeted by HIV-1 Vpu^10^) is upregulated by T cell activation, then progressively depleted in HIV-infected (red, SBP-ΔLNGFR positive) but not uninfected (blue, SBP-ΔLNGFR negative) cells (**Fig. 2c, left panel**). Conversely, the restriction factor SAMHD1 (targeted by some HIV-2/SIV Vpx and Vpr variants, but not HIV-1^23^–^25^) is depleted by T cell activation, independent of HIV infection (**Fig. 2c, right panel**). In these graphical representations, relative protein abundances for each condition are depicted by bars, and ratios of protein abundances in paired experimental/control cells from each condition/time point are depicted by lines (grey, resting/activated; red, SBP-ΔLNGFR positive, infected; blue, SBP-ΔLNGFR negative, uninfected).

Aside from tetherin, levels of many other reported Vpu (CD4, SNAT1)^2,19^, Nef (CD4, SERINC5)^11,12,18^, Vif (APOBEC3 and PPP2R5 families)^3,9^ and Vpr (UNG, HLTF, ZGPAT, VPRBP, MUS81, EME1, MCM10, TET2)^26–33^ substrates were all reduced by HIV infection in primary T cells (**Fig. 2d, and Supplementary Fig. 3b**). Conversely, and consistent with our previous observations in CEM-T4s, APOBEC3B and SERINC1 were not depleted (**Supplementary Fig. 3b**)^2,3^. In the absence of donor haplotyping, polymorphisms at the MHC-I locus make routine proteomic quantification problematic. Nonetheless, our data are consistent with depletion of HLA-A and HLA-B, but not HLA-C (**Supplementary Fig. 3b**), as previously reported for Nef/Vpu variants from NL4-3 HIV^34–36^.

Together with cellular proteins, we identified gene products from 7 viral open reading frames (ORFs; **Supplementary Fig. 3c**). As expected^37^, viral regulatory and accessory proteins expressed from fully spliced, Rev-independent transcripts (Tat, Rev, Nef-P2A and SBP-ΔLNGFR) were expressed early in infection, peaking at 24 hrs. Conversely, viral structural proteins expressed from unspliced, Rev-dependent transcripts (Gag and Gagpol) were expressed late in infection, increasing progressively from 24 to 48 hrs. Viral accessory proteins expressed from partially spliced transcripts were either not detected (Vpr and Vpu) or incompletely quantitated (Vif).

Inter-individual variability is known to affect gene expression during T cell activation^38^. Accordingly, to identify reproducible HIV targets, we analysed primary human CD4+ T cells from 3 further donors. In each case, mock-infected cells were compared with HIV-infected cells selected using AFMACS 48 hrs post-infection (**Fig. 3a,b and Supplementary Fig. 4a**). Aside from APOBEC3 proteins, we recently discovered the PPP2R5A-E (B56) family of PP2A phosphatase regulatory subunits to be degraded by diverse Vif variants, spanning primate and ruminant lentiviruses^3^. To formally document Vif-dependent changes in primary T cells, both wildtype (WT) and Vif-deficient (ΔVif) viruses were therefore included. Whilst some donor-dependent differences were apparent, most sample-sample variability was accounted for by HIV infection (**Supplementary Fig. 4b**), and all accessory protein substrates from **Fig. 2d and Supplementary Fig. 3b** were significantly depleted by WT HIV (**Fig. 3c – left panel**). In total, we quantitated 8,789 cellular proteins, of which 650 were significantly regulated by HIV infection (q<0.05).

**Figure 3:**
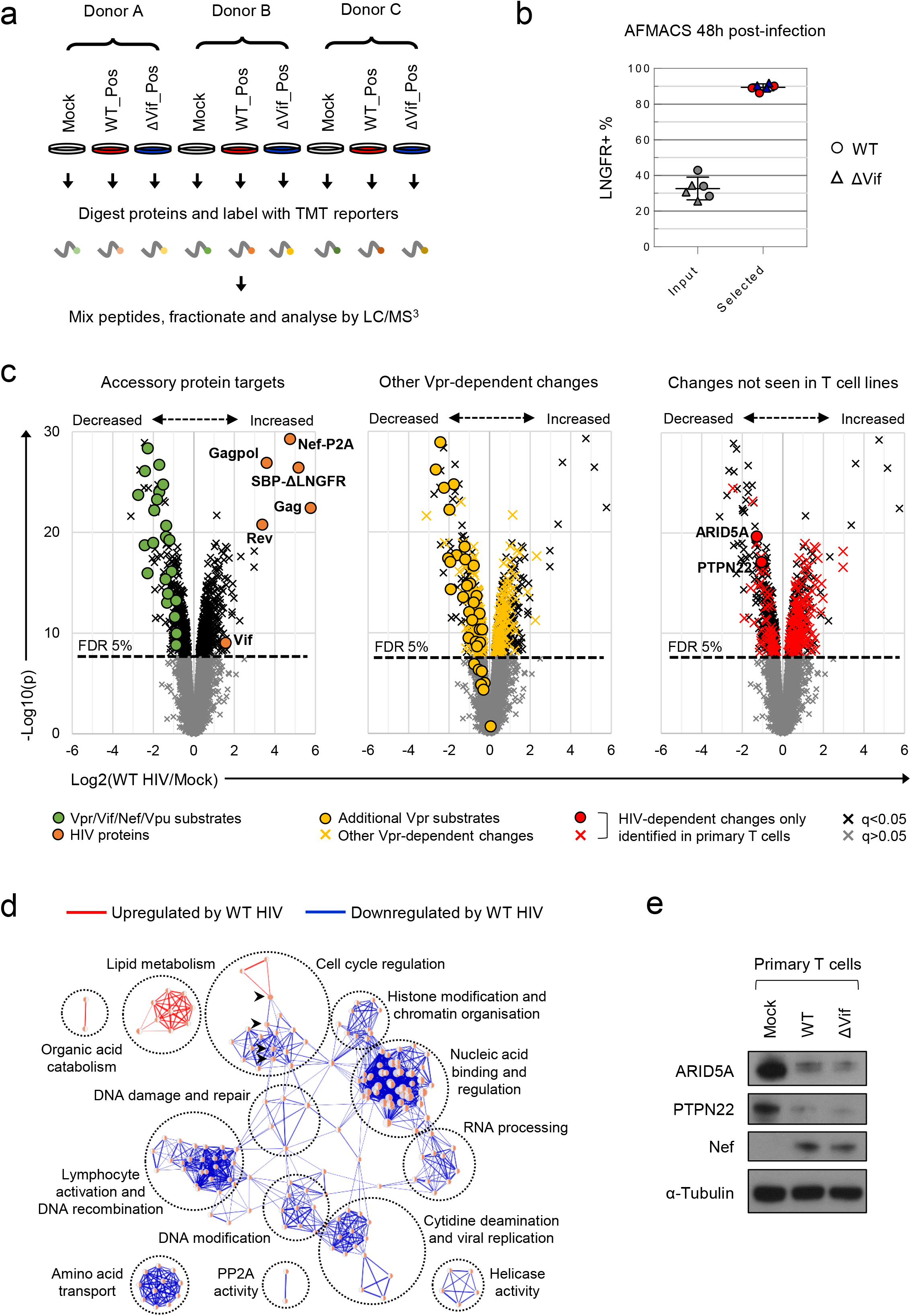
Proteins and pathways regulated by HIV in primary T cells. **a**, Overview of single time point proteomic experiment. HIV-infected (LNGFR+) primary T cells were isolated using AFMACS 48 hrs post-infection with WT (red) or ΔVif (blue) HIV-AFMACS. **b**, AFMACS-based enrichment of WT (red circles) and ΔVif (blue triangles) HIV-infected (LNGFR+) cells used for **a**, with means and 95% CIs. Corresponding cells pre-selection are included for each donor/virus (WT, grey circles; ΔVif, grey triangles). Cells were stained with anti-LNGFR and anti-CD4 antibodies and analysed by flow cytometry, with representative data in **Supplementary Fig. 4a**. **c**, Protein abundances in WT HIV-infected vs mock-infected cells from **a**. Volcano plots show statistical significance (y axis) vs fold change (x axis) for 8,795 cellular and viral proteins quantitated in cells from all 3 donors (no missing values). Proteins with Benjamini-Hochberg false discovery rate (FDR)-adjusted p values (q values) <0.05 or >0.05 are indicated (FDR threshold of 5%). Proteins highlighted in each plot are summarised in the legend. Vpr/Vif/Nef/Vpu substrates (green circles) comprise proteins from **Fig. 2c,d** and **Supplementary Fig. 3b**, excluding negative controls (SAMHD1, APOBEC3B, SERINC1, HLA-C) and HLA-A/B alleles (different in each donor) but including SMUG1 (not identified in time course proteomic experiment)^26^ and both quantitated isoforms of PP2R5C (Q13362 and Q13362-4) and ZGPAT (Q8N5A5 and Q8N5A5-2). Additional Vpr substrates (gold circles) and Vpr-dependent changes (gold crosses) comprise recently described direct and indirect Vpr targets^39^. HIV-dependent changes only identified in primary T cells (red circles and crosses) comprise proteins with q<0.05 either not identified or not concordantly regulated by HIV in CEM-T4s^3^ (and exclude known accessory protein-dependent changes). Further details on comparator datasets used in this figure are provided in the **Methods**. **d**, Enrichment Map^85^ network-based visualisation of Gene Ontology (GO) functional annotation terms enriched amongst upregulated (red) or downregulated (blue) proteins with q<0.05 in WT HIV-infected vs mock-infected cells from **a**. Each node represents a GO term, with node size indicating number of annotated proteins, edge thickness representing degree of overlap, and similar GO terms placed close together. Degree of enrichment is mapped to node colour (left side, enriched amongst upregulated proteins; right side, enriched amongst downregulated proteins) as a gradient from white (no enrichment) to red (high enrichment). Highlighted nodes (arrow heads) represent GO terms enriched amongst both upregulated and downregulated proteins. **e**, Depletion of ARID5A and PTPN22 by HIV. AFMACS-selected (LNGFR+) HIV-infected cells from **a** (donor A) were lysed in 2% SDS and analysed by immunoblot with anti-ARID5A, anti-PTPN22, anti-Nef and anti-α-tubulin antibodies. Same lysates as **Fig. 4d**.

Compared with a previous, similar experiment using CEM-T4s^3^, we observed greater variability in protein abundances between replicates (**Supplementary Fig. 5a**), but a high degree of correlation in HIV-dependent changes between cell types (**Supplementary Fig. 5b**). As well as “canonical” accessory protein targets, we have recently discovered that most protein-level changes in HIV-infected CEM-T4s may be explained by primary and secondary effects of Vpr, including degradation of at least 34 additional substrates^39^. These changes were recapitulated in primary T cells (**Fig. 3c – middle panel**), with 33 newly described Vpr substrates quantitated, and 32 decreased in abundance. Several other cell surface proteins reported to be downregulated by Nef and/or Vpu were also depleted, but the magnitude of effect was typically modest, and many were unchanged (**Supplementary Fig. 5c – left panel**). Likewise, we did not see evidence of HIV/Vif-dependent transcriptional regulation of RUNX1 target gene products such as T-bet/TBX21 (**Supplementary Fig. 5c – middle panel**)^40^. Nonetheless, taken together, known accessory protein-dependent changes, characterised in transformed T cell lines, are able to account for 297/650 (46%) of proteins regulated by HIV in primary T cells (**Supplementary Fig. 5d**), including 175/299 (59%) of proteins decreased in abundance.

As with individual proteins, pathways and processes downregulated by HIV infection of primary T cells are dominated by the effects of accessory proteins (**Fig. 3d and Supplementary Fig. 6a**). These include the DNA damage response and cell cycle (Vpr)^39,41–47^, cytidine deamination and PP2A activity (Vif)^3,9,48^ and amino acid transport (Vpu/Nef)^2^. Proteins upregulated by HIV are more diverse, with fewer dominant functional clusters. Nonetheless, we saw marked increases in proteins associated with lipid and sterol metabolism (**Supplementary Fig. 6b**). A similar effect has been reported in T cell lines at the transcriptional level, and attributed to the expression of Nef^49,50^. Similarly, several proteins in these pathways are indirectly regulated by Vpr (**Supplementary Fig. 6b**)^39^.

Despite the overall agreement with cell line data, 1,252/8,789 (14%) of cellular proteins quantitated in this experiment were not identified in a previous, similar experiment using CEM-T4s^3^. Furthermore, having excluded known accessory-protein dependent changes, 192/650 (30%) proteins regulated by HIV in primary T cells are either not detected, or not significantly/concordantly regulated, in CEM-T4s (**Fig. 3c – right panel and Supplementary Fig. 5d**). These proteins may represent accessory protein substrates expressed in primary T cells but not T cell lines, or proteins regulated by alternative, cell type-specific mechanisms, such as the interferon response (**Supplementary Fig. 5c – right panel**)^51^. To validate our proteomic data, we focused on 2 such novel HIV targets with commercially available antibodies: ARID5A and PTPN22. These proteins were readily identified in proteomic datasets from primary T cells (9-14 unique peptides) but not CEM-T4s, and consistently depleted across all donors with a fold change >2 (**Supplementary Fig. 7a**). As expected, depletion was also seen by immunoblot (**Fig. 3e**).

We previously showed that substrates of different HIV accessory proteins could be distinguished by their characteristic patterns of temporal regulation in HIV-infected CEM-T4s^3^, and similar clustering was observed in primary T cells (**Supplementary Fig. 7b**). Vpr is packaged stoichiometrically in virions^52–54^ and, since the number of fusogenic HIV particles exceeds the infectious MOI by at least several fold^55^, all cells in our time course experiment were necessarily exposed to incoming Vpr. Accordingly, depletion of known Vpr substrates was near-maximal by 24 hrs in infected (red, SBP-ΔLNGFR positive) cells, with partial depletion also seen in uninfected (blue, SBP-ΔLNGFR negative) cells (**Fig. 2e – upper panel**). In contrast, since *de novo* viral protein synthesis is absolutely required, depletion of known Vif, Nef and Vpu substrates increased progressively from 24 to 48 hrs, and was only seen in HIV-infected (red, SBP-ΔLNGFR positive) cells (**Fig. 2e – lower panel**). Based on their patterns of temporal regulation, ARID5A and PTPN22 are therefore very likely to represent novel Vpr substrates, specific for primary T cells (**Supplementary Fig. 7c**). Consistent with this, another member of the ARID5 subfamily of AT-rich interaction domain (ARID)-containing proteins, ARID5B, is a widely conserved target of Vpr variants from primate lentivuses^39^, and shares a similar temporal profile (**Supplementary Fig. 7d**).

In respect of Vif targets, as predicted, both APOBEC3 and PPP2R5 family proteins were depleted in primary CD4+ T cells infected with WT, but not ΔVif viruses (**Fig. 4a,b**). Vif-dependent depletion of PPP2R5A-E causes a marked increase in protein phosphorylation in HIV-infected CEM T4 T cells, particularly substrates of the aurora kinases (AURKA/B)^3^. AURKB activity is enhanced by “activation loop” auto-phosphorylation at threonine 232 (T232), antagonised by PP2A-B56^56,57^. Accordingly, a marked increase in AURKB T232 phosphorylation is seen in CEM-T4s transduced with Vif as a single transgene (**Fig. 4c**). We therefore confirmed depletion of PPP2R5D by immunoblot of AFMACS-selected primary T cells and, as a functional correlate, observed increased AURKB phosphorylation (**Fig. 4d**).

**Figure 4:**
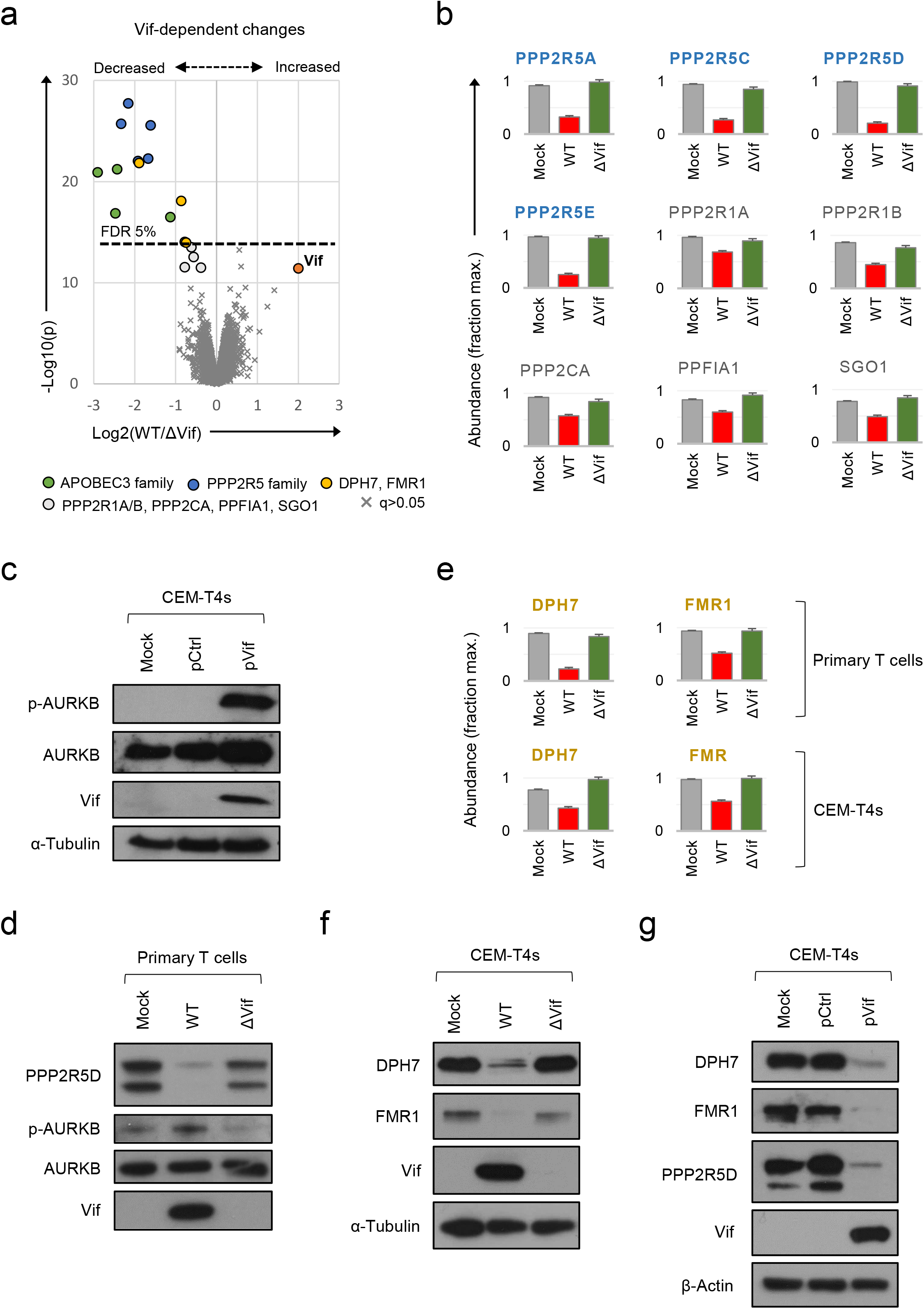
Vif-dependent cellular targets in primary T cells. **a**, Protein abundances in WT HIV-infected vs ΔVif HIV-infected cells from single time point proteomic experiment (**Fig. 3a**). Statistical significance (y axis) vs fold change (x axis) is shown for 8,795 cellular and viral proteins quantitated in cells from all 3 donors (no missing values). Proteins with Benjamini-Hochberg false discovery rate (FDR)-adjusted p values (q values) <0.05 or >0.05 are indicated (FDR threshold of 5%). Highlighted groups of differentially regulated proteins are summarised in the legend, including 2 quantitated isoforms of PP2R5C (Q13362 and Q13362-4) and FMR1 (Q06787 and Q06787-2). **b**, Abundances of Vif-dependent PPP2R5 family and related proteins highlighted in **a** in mock-infected (grey), WT HIV-infected (red) and ΔVif HIV-infected (green) cells from single time point proteomic experiment (**Fig. 3a**). Mean abundances (fraction of maximum) with 95% CIs are shown. Only the canonical isoform of PPP2R5C (Q13362) is shown. **c**, Activation of AURKB by Vif. CEM-T4s were mock-transduced or transduced with control (pCtrl) or Vif-expressing (pVif) lentiviruses for 48 hrs (62-78% GFP+), lysed in 2% SDS and analysed by immunoblot with anti-phospho-AURK (T232), anti-total AURKB, anti-Vif and anti-α-tubulin antibodies. **d**, Vif-dependent activation of AURKB. AFMACS-selected (LNGFR+) HIV-infected cells from **Fig. 3a** (donor A) were lysed in 2% SDS and analysed by immunoblot with anti-PPP2R5D, anti-phospho-AURK (T232), anti-total AURKB and anti-Vif antibodies. Same lysates as **Fig. 3e** **e**, Abundances of DPH7 and FMR1 in mock-infected (grey), WT HIV-infected (red) and ΔVif HIV-infected (green) primary T cells from single time point proteomic experiment (**Fig. 3a**) and CEM-T4s^3^. Mean abundances (fraction of maximum) with 95% CIs intervals are shown. Only the canonical isoform of FMR1 (Q06787) is shown. **f**, Vif-dependent depletion of DPH7 and FMR1. CEM-T4s were mock-infected or infected with WT or ΔVif HIV-AFMACS viruses for 48 hrs (77-82% LNGFR+ cells), lysed in 2% SDS, and analysed by immunoblot with anti-DPH7, ant-FMR1, anti-Vif and anti-α-tubulin antibodies. **g**, Depletion of DPH7, FMR1 and PPP2R5D by Vif. CEM-T4s were mock-transduced or transduced with control (pCtrl) or Vif-expressing (pVif) lentiviruses for 48 hrs (86-88% GFP+ cells), lysed in 2% SDS and analysed by immunoblot with anti-phospho-AURK, anti-total AURKB, anti-Vif and anti-α-tubulin (loading control) antibodies.

Besides these known substrates, we also noted differential regulation of several other proteins in primary T cells infected with WT vs ΔVif viruses (**Fig. 4a**). Modest changes in PPP2R1A, PPP2R1B and PPP2CA (catalytic/structural subunits of the trimeric PP2A holoenzyme) and PPFIA1 and SGO1 (known PP2A interactors)^58–60^ are likely to be secondary to destabilisation of PP2A by PPP2R5 depletion, or reflect proximity of the holoenzyme to the Vif-cullin E3 ligase complex (**Fig. 4b**). Conversely, DPH7 and FMR1 are not known to interact physically with PP2A, and show more profound and consistent depletion (**Fig. 4e**). We therefore suspected these proteins to be novel Vif substrates.

To confirm these findings, we first re-examined our proteomic data from CEM-T4s^3^. Unlike ARID5A and PTPN22, DPH7 and FMR1 are expressed in CEM-T4 as well as primary T cells, and only decreased in abundance in HIV-infected cells in the presence of Vif (**Fig. 4e**). Next, we confirmed Vif-dependent depletion of both proteins by immunoblot, in cells infected with WT (but not ΔVif) viruses (**Fig. 4f**). Finally, we repeated these observations in cells transduced with Vif as a single transgene (**Fig. 4g**). Vif is therefore both necessary and sufficient for depletion of DPH7 and FMR1 and, taken together with APOBEC3 and PPP2R5 family proteins/interactors, we can account for all significant Vif-dependent changes in the natural target cell of HIV infection.

Compared with FACS, bead-based magnetic sorting is fast, simple and scalable for simultaneous processing of multiple samples and large cell numbers^61^. In conventional, antibody-based immunomagnetic selection, cells remain coated with beads and antibody-antigen complexes, risking alteration of their behaviour or viability through cross-linking of cell-surface receptors or internalisation of ferrous beads^61–63^. Conversely, AFMACS-based selection is antibody free, and selected cells are released from the beads by incubation with biotin, suitable for a full range of downstream applications^14^. The HIV-AFMACS virus described in this study allows routine isolation of HIV-infected cells subjected to an MOI<1, avoiding artefacts associated with high MOIs and facilitating experiments in primary cells, where high levels of infection are difficult to achieve in practice.

To demonstrate the utility of this system and provide a resource for the community, we have generated the first high coverage proteomic atlas of HIV-infected primary human CD4+ T cells. As well as identifying HIV-dependent changes in cells from multiple donors, viral regulation may be assessed against a background of endogenous regulation triggered by T cell activation. Unlike T cell lines, primary T cells express a full range of proteins relevant to HIV infection *in vivo*, and are not confounded by the genetic and epigenetic effects of transformation. Furthermore, proteins unique to primary T cells were significantly more likely to be regulated by HIV infection than proteins detected in both primary T cells and CEM-T4s (131/1,252=10.5% vs 519/7,537=6.9%; p<0.0001, two-tailed Fisher’s exact test). Our data validate many, but not all, previously reported HIV accessory protein targets. For some Vpu/Nef substrates, such as NTB-A and CCR7, downregulation from the plasma membrane may occur in the absence of protein degradation^64–66^. For others, such as ICAM-1/3, accessory protein expression may prevent upregulation, without reducing levels below baseline^67^. Alternatively, and particularly where targets have been discovered using model cell line or overexpression systems, regulation may not be quantitatively significant at the protein level in the context and/or natural cell of HIV infection.

Previous temporal proteomic analyses of HIV-infected primary human CD4+ T cells were hampered by limited coverage of the cellular proteome (<2,000 proteins), and did not detect regulation of known or novel HIV accessory protein targets^68,69^. A more recent study quantitated 7,761 proteins in FACS-sorted T cells at a single time point 96 hrs post-infection with an R5-tropic, GFP-expressing Nef-deficient virus^70^. Depletion of several accessory protein targets (including APOBEC3 and PPP2R5 families) was confirmed, and many proteins differentially regulated at 96 hrs were also regulated at 48 hrs in our study (**Supplementary Fig. 8a**). In keeping with the late time point, changes were dominated by pathways involved in cell death and survival, and factors maintaining viability of HIV-infected T cells (such as BIRC5) were enriched. Conversely, the full dataset is not available, effect sizes were generally compressed (**Supplementary Fig. 8a**), and 413/650 (64%) of the HIV-dependent changes identified in our study were obscured, including depletion of ARID5A, PTPN22, DPH7 and 19/51 known accessory protein substrates (**Supplementary Fig. 8b**).

To maximise viral titers, enable synchronous single round infections and avoid co-receptor-dependent targeting of T cell lineages with pre-existing proteomic differences, we used an Env-deficient proviral backbone and pseudotyped viruses with VSVg. Different envelope proteins may alter the route of viral entry, and/or Env-dependent signalling early in infection^71,72^. To limit the impact of these effects, we focussed on cellular proteins progressively regulated over 48 hrs infection, and included SBP-ΔLNGFR negative cells (flow-through, exposed to incoming virions but not infected) as a control in our time course analysis. Temporal profiling is particularly well suited to identifying and characterising host factors regulated directly by viral proteins. In fact, as we show here, HIV accessory proteins account for much or most of the proteomic remodelling in infected cells. The abundance of direct accessory protein targets likely explains why proteins and processes/pathways downregulated by HIV in primary T cells correlate so well with changes seen in T cell lines, and are robust to inter-individual variation. In comparison, upregulated proteins concord less well with changes in T cell lines, and functional effects are less homogeneous. This may be because upregulated proteins reflect indirect effects (for example, secondary changes in transcription), which are more likely to be cell-type specific.

Amongst the HIV accessory proteins, Vif was thought until recently to exclusively degrade APOBEC3 family cytidine deaminases. As well as confirming equivalently-potent Vif-dependent depletion of PPP2R5 family phosphatase subunits in primary T cells, our data revealed unexpected Vif-dependent depletion of DPH7 and FMR1. Further work will be required to confirm that these proteins are recruited directly for degradation by the ubiquitin-proteasome system, determine whether (like APOBEC3 and PPP2R5 proteins) they are antagonised by Vif variants from diverse primate and non-primate lentiviral lineages, and identify relevant *in vitro* virological phenotypes associated with target depletion. Nonetheless, FMR1 is known to reduce HIV virus infectivity when over-expressed in producer cells^73^, and the other novel targets highlighted in this study also impact processes relevant for HIV replication, such as inflammatory cytokine signalling (ARID5A)^74^, T cell activation (PTPN22)^75^ and translational fidelity (DPH7)^76,77^. The diversity of these targets underscores the benefit of an unbiased, systems-level approach to viral infection, and the capacity of the resources presented in this study to reveal unsuspected aspects of the host-virus interaction in the natural target cell of HIV infection.

## Methods

### General cell culture

CEM-T4 T cells (CEM-T4s)^5^ were obtained from the AIDS Reagent Program, Division of AIDS, NIAD, NIH: Dr J.P. Jacobs and cultured in RPMI supplemented with 10 % FCS, 100units/ml penicillin and 0.1 mg/ml streptomycin at 37 °C in 5 % CO2. HEK 293T cells (Lehner laboratory stocks) were cultured in DMEM supplemented with 10 % FCS, 100units/ml penicillin and 0.1 mg/ml streptomycin at 37 °C in 5 % CO2. All cells were confirmed to be mycoplasma negative (Lonza MycoAlert). Cell line authentication was not undertaken.

### Primary cell isolation and culture

Primary human CD4+ T cells were isolated from peripheral blood by density gradient centrifugation over Lympholyte-H (Cedarlane Laboratories) and negative selection using the Dynabeads Untouched Human CD4 T Cells kit (Invitrogen) according to the manufacturer’s instructions. Purity was assessed by flow cytometry for CD3 and CD4 and routinely found to be ≥95%. Cells were activated using Dynabeads Human T-Activator CD3/CD28 beads (Invitrogen) according to the manufacturer’s instructions and cultured in RPMI supplemented with 10% FCS, 30U/ml recombinant human IL-2 (PeproTech), 100units/ml penicillin and 0.1mg/ml streptomycin at 37°C in 5% CO2.

### Ethics statement

Ethical permission for this study was granted by the University of Cambridge Human Biology Research Ethics Committee (HBREC.2017.20). Written informed consent was obtained from all volunteers prior to providing blood samples.

### HIV-1 molecular clones

pNL4-3-ΔEnv-EGFP^7^ was obtained from the AIDS Reagent Program, Division of AIDS, NIAD, NIH: Drs Haili Zhang, Yan Zhou, and Robert Siliciano and the complete proviral sequence verified by Sanger sequencing (Source BioScience). Derived from the HIV-1 molecular clone pNL4-3, it encodes EGFP in the *env* ORF), resulting in a large, critical *env* deletion and expression of a truncated Env-EGFP fusion protein retained in the endoplasmic reticulum (ER) by a 3’ KDEL ER-retention signal.

The SBP-ΔLNGFR selection marker comprises the high-affinity 38 amino acid SBP fused to the N-terminus of a truncated (non-functional) member of the Tumour Necrosis Factor Receptor superfamily (LNGFR)^14^. As a type I transmembrane glycoprotein, expression at the cell surface requires a 5’ signal peptide.

To replace EGFP with SBP-ΔLNGFR (generating pNL4-3-ΔEnv-SBP-ΔLNGFR) a synthetic gene fragment (gBlock; Integrated DNA Technologies, IDT) was incorporated into pNL4-3-ΔEnv-EGFP by Gibson assembly between SalI/BsaBI sites (**gBlock #1; Supplementary Methods**). In this construct, SBP-ΔLNGFR is fused with the endogenous Env signal peptide.

To express SBP-ΔLNGFR downstream of *nef* and a “self-cleaving” *Porcine teschovirus-1* 2A (P2A) peptide (generating pNL4-3-ΔEnv-EGFP-Nef-P2A-SBP-ΔLNGFR) a gBlock (IDT) was incorporated into pNL4-3-ΔEnv-EGFP by Gibson assembly between HpaI/XhoI sites (**gBlock #2; Supplementary Methods**). In this construct, SBP-ΔLNGFR is co-translated with codon-optimised Nef (Nef-hu) and includes an exogenous murine immunoglobulin (Ig) signal peptide. SBP-ΔLNGFR was located downstream (rather than upstream) of Nef-hu to avoid disruption of Nef myristoylation by addition of a 5’ proline residue following P2A “cleavage”.

To express SBP-ΔLNGFR downstream of *nef* and an *Encephalomyocarditis virus* (EMCV) internal ribosome entry site (IRES; generating pNL4-3-ΔEnv-EGFP-Nef-IRES-SBP-ΔLNGFR) a gBlock (IDT) was incorporated into pNL4-3-ΔEnv-EGFP by Gibson assembly between HpaI/XhoI sites (**gBlock #3; Supplementary Methods**). In this construct, SBP-ΔLNGFR is translated independently of Nef-hu and includes an exogenous murine Ig signal peptide. A widely-used replication-competent HIV EGFP reporter virus was previously generated using a similar approach^78,79^.

In all constructs, Nef or Nef-hu expression is mediated by the WT HIV LTR promoter and naturally occurring splice sites. In constructs with a P2A peptide or IRES, the use of codon-optimised Nef-hu minimises homology with the U3 region of the 3’ LTR (overlapped by the endogenous *nef* sequence) and reduces the risk of recombination.

To remove EGFP from constructs with a P2A peptide or IRES, a gBlock (IDT) was incorporated by Gibson assembly between SalI/BsaBI sites (**gBlock #4; Supplementary Methods**). To avoid generating a truncated protein product fused to the Env signal peptide, the *env* start codon and other potential out of frame start codons in the *vpu* ORF were disrupted with point mutations, whilst maintaining the Vpu protein sequence. Redundant *env* sequence was minimised, without disrupting the Rev response element (RRE).

To truncate the U3 region of the 3’ LTR in constructs with a P2A peptide or IRES (with or without EGFP), a gBlock (IDT) was incorporated by Gibson assembly between XhoI/NaeI sites (**gBlock #5; Supplementary Methods**). Previous studies have shown that, in the presence of an intact *nef* ORF, the overlap between *nef* and the U3 region is dispensable for HIV gene expression and replication^80^.

To generate a Vif-deficient HIV-AFMACS molecular clone (pNL4-3-ΔVif-ΔEnv-Nef-P2A-SBP-ΔLNGFR), an AgeI/PflMI restriction fragment was subcloned from pNL4-3-ΔVif-ΔEnv-EGFP^3^ into pNL4-3-ΔEnv-Nef-P2A-SBP-ΔLNGFR (HIV-AFMACS).

Where appropriate, additional unique restriction sites were included to facilitate future cloning. All sequences were verified by Sanger sequencing (Source BioScience). The complete HIV-AFMACS (pNL4-3-ΔEnv-Nef-P2A-SBP-ΔLNGFR) sequence is available in the **Supplementary Methods**.

### Lentivectors for transgene expression

For over-expression of codon optimised NL4-3 Vif (Vif-hu) in CEM-T4s, a gBlock (IDT) was incorporated into pHRSIN-SE-P2A-SBP-ΔLNGFR-W^14^ in place of SBP-ΔLNGFR by Gibson assembly between KpnI/XhoI sites to generate pHRSIN-SE-P2A-Vif-hu-W (in which Vif-hu and EGFP expression are coupled by a P2A peptide).

### Viral stocks

VSVg-pseudotyped NL4-3-ΔEnv-based viral stocks were generated by co-transfection of 293T cells with pNL4-3-ΔEnv molecular clones and pMD.G at a ratio of 9:1 (μg) DNA and a DNA:FuGENE 6 ratio of 1 μg:6 μl. Media was changed the next day and viral supernatants harvested and filtered (0.45 μm) at 48 h prior to concentration with LentiX Concentrator (Clontech) and storage at −80 °C.

VSVg-pseudotyped pHRSIN-based lentivector stocks were generated by co-transfection of 293Ts with lentivector, p8.91 and pMD.G at a ratio of 2:1:1 (μg) DNA and a DNA:FuGENE 6 ratio of 1ug:3μl. Viral supernatants were harvested, filtered, concentrated and stored as per NL4-3-ΔEnv-based viral stocks.

All viruses were titered by infection/transduction of known numbers of relevant target cells under standard experimental conditions followed by flow cytometry for LNGFR and/or CD4/EGFP at 48 h to identify % infected cells.

### T cell infections

CEM-T4s were infected/transduced by spinoculation at 800 g for 2 h in a non-refrigerated benchtop centrifuge in complete media supplemented with 10mM HEPES. Primary human CD4+ T cells were infected using the same protocol 48 h after activation with CD3/CD28 Dynabeads.

### Antibody-Free Magnetic Cell Sorting (AFMACS)

AFMACS-based selection of CEM-T4 or primary human CD4+ T cells using the streptavidin-binding SBP-ΔLNGFR affinity tag was carried out essentially as previously described^14^. For primary T cells, CD3/CD28 Dynabeads were first removed using a DynaMag-2 magnet (Invitrogen). 24 or 48 h post-infection, washed cells were resuspended in incubation buffer (IB; Hank’s balanced salt solution, 2% dialysed FCS, 1x RPMI Amino Acids Solution (Sigma), 2 mM L-glutamine, 2mM EDTA and 10mM HEPES) at 10e7 cells/ml and incubated with Biotin Binder Dynabeads (Invitrogen) at a bead-to-total cell ratio of 4:1 for 30 mins at 4oC. Bead-bound cells expressing SBP-ΔLNGFR were selected using a DynaMag-2 (Invitrogen), washed to remove uninfected cells, then released from the beads by incubation in complete RPMI with 2 mM biotin for 15 mins at room temperature (RT). Enrichment was routinely assessed by flow cytometry pre- and post-selection.

### Proteomic analysis

#### Sample preparation

For TMT-based whole cell proteomic analysis of primary human CD4+ T cells, resting or activated cells were washed with ice-cold PBS with Ca/Mg pH 7.4 (Sigma) and frozen at −80 °C. Samples were lysed, reduced, alkylated, digested and labelled with TMT reagents using the iST-NHS Sample Preparation Kit (PreOmics GmbH) according to the manufacturer’s instructions. Typically, 5e6 resting or 1e6 activated cells were used for each condition. Where indicated, cells were spinoculated with VSVg-pseudotyped viruses at an effective multiplicity of infection (MOI) of approximately 0.5 and subjected to AFMACS-based selection 24 or 48 h post-infection.

#### Off-line high pH reversed-phase (HpRP) peptide fractionation

HpRP fractionation was conducted on an Ultimate 3000 UHPLC system (Thermo Scientific) equipped with a 2.1 mm × 15 cm, 1.7 μm Acquity BEH C18 column (Waters, UK). Solvent A was 3% ACN, solvent B was 100% ACN, and solvent C was 200 mM ammonium formate (pH 10). Throughout the analysis C was kept at a constant 10%. The flow rate was 400 μL/min and UV was monitored at 280 nm. Samples were loaded in 90% A for 10 min before a gradient elution of 0–10% B over 10 min (curve 3), 10-34% B over 21 min (curve 5), 34-50% B over 5 min (curve 5) followed by a 10 min wash with 90% B. 15 s (100 μL) fractions were collected throughout the run. Peptide-containing fractions were orthogonally recombined into 24 fractions (e.g. fractions 1, 25, 49, 73 and 97) and dried in a vacuum centrifuge. Fractions were stored at −80 °C prior to analysis.

#### Mass spectrometry

Data were acquired on an Orbitrap Fusion mass spectrometer (Thermo Scientific) coupled to an Ultimate 3000 RSLC nano UHPLC (Thermo Scientific). HpRP fractions were resuspended in 20 μl 5% DMSO 0.5% TFA and 10 uL injected. Fractions were loaded at 10 μl/min for 5 min on to an Acclaim PepMap C18 cartridge trap column (300 um × 5 mm, 5 um particle size) in 0.1% TFA. Solvent A was 0.1% FA and solvent B was ACN/0.1% FA. After loading, a linear gradient of 3–32% B over 3 h was used for sample separation over a column of the same stationary phase (75 μm × 50 cm, 2 μm particle size) before washing at 90% B and re-equilibration.

An SPS/MS3 acquisition was used for all samples and was run as follows. MS1: Quadrupole isolation, 120’000 resolution, 5e5 AGC target, 50 ms maximum injection time, ions injected for all parallelisable time. MS2: Quadrupole isolation at an isolation width of m/z 0.7, CID fragmentation (NCE 35) with the ion trap scanning out in rapid mode from m/z 120, 8e3 AGC target, 70 ms maximum injection time, ions accumulated for all parallelisable time. In synchronous precursor selection mode the top 10 MS2 ions were selected for HCD fragmentation (65NCE) and scanned out in the orbitrap at 50’000 resolution with an AGC target of 2e4 and a maximum accumulation time of 120 ms, ions were not accumulated for all parallelisable time. The entire MS/MS/MS cycle had a target time of 3 s. Dynamic exclusion was set to +/−10 ppm for 90 s, MS2 fragmentation was trigged on precursor ions 5e3 counts and above.

#### Data processing and analysis

Spectra were searched by Mascot within Proteome Discoverer 2.1 in two rounds. The first search was against the UniProt Human reference proteome (26/09/17), the HIV-AFMACS proteome and compendium of common contaminants (GPM). The second search took all unmatched spectra from the first search and searched against the human trEMBL database (Uniprot, 26/09/17). The following search parameters were used. MS1 Tol: 10 ppm. MS2 Tol: 0.6 Da. Fixed Mods: Ist-alkylation (+113.084064 Da) (C) and TMT (N-term, K). Var Mods: Oxidation (M). Enzyme: Trypsin (/P). MS3 spectra were used for reporter ion-based quantitation with a most confident centroid tolerance of 20 ppm. PSM FDR was calculated using Mascot percolator and was controlled at 0.01% for ‘high’ confidence PSMs and 0.05% for ‘medium’ confidence PSMs. Normalisation was automated and based on total s/n in each channel. Proteins/peptides satisfying at least a ‘medium’ FDR confidence were taken forth to statistical analysis in R. This consisted of a moderated T-test (Limma) with Benjamini-Hochberg correction for multiple hypotheses to provide a q value for each comparison^81^. Further data manipulation and general statistical analysis was conducted using Excel, XLSTAT and GraphPad Prism 7.

For functional analysis of proteins significantly downregulated or upregulated by WT HIV (q<0.05) in the single time point proteomic experiment (**Fig. 3a**), enrichment of Gene Ontology (GO) Biological Process (GOTERM_BP_FAT) and Molecular Function (GOTERM_MF_FAT) terms against a background of all proteins quantitated was determined using the Database for Annotation, Visualization and Integrated Discovery (DAVID) 6.8 (accessed on 22/7/2018 at https://david.ncifcrf.gov/) with default settings^82,83^. Human proteins annotated to GO:0016126 (sterol biosynthetic process) were retrieved from AmiGO 2 (accessed on 27/7/2018 at http://amigo.geneontology.org/amigo)^84^. To account for redundancy between annotations, enriched GO terms were visualised using the Enrichment Map 3.1.0 plugin^85^ for Cytoscape 3.6.1. (downloaded from http://cytoscape.org/)^86^ with default settings (q value cut-off of 0.1) and sparse-intermediate connectivity. Clusters were manually labelled to highlight the prevalent biological functions amongst each set of related annotations.

For clustering according to profiles of temporal expression, known accessory protein substrates from Fig. 2c,d and Supplementary Fig. 3b and additional Vpr substrates shown in Fig. 3c were analysed using Cluster 3.0 (downloaded from http://bonsai.hgc.jp/~mdehoon/software/cluster/software.htm)^87^ and visualised using Java TreeView 1.1.6r4 (downloaded from http://jtreeview.sourceforge.net)^88^. Only proteins significantly downregulated by WT HIV (q<0.05) in the single time point proteomic experiment (**Fig. 3a**) were included. Where more than one isoform was quantitated, only the canonical isoform was used (PPP2R5C, Q13362; ZGPAT, Q8N5A5; NUSAP1, Q9BXS6). Data from the time course proteomic experiment (**Fig. 2a**) were expressed as log2(ratio)s of abundances in experimental (Expt):control (Ctrl) cells for each condition/time point, and range-scaled to highlight patterns of temporal expression relative to the biological response range (minimum-maximum) for each protein. Agglomerative hierarchical clustering was performed using uncentered Pearson correlation and centroid linkage^89^.

#### Comparison with CEM-T4 T cells

To compare results between primary and transformed T cells at a similar depth of proteomic coverage, we re-analysed TMT-labelled peptide eluates from a previous study^3^ conducted in CEM-T4s spinoculated in triplicate with VSVg-pseudotyped NL4-3-ΔEnv-EGFP WT or ΔVif viruses at an MOI of 1.5. This extended analysis consisted of reinjection of HpRP fractions on longer (3 h) gradients using a higher performance (75 as opposed to 50 cm) analytical column and the MS parameters employed in this study. In total, the new CEM-T4 dataset covered 8,065 proteins, comparable with the datasets from primary T cells described here.

#### Comparisons with other published datasets

We have recently characterised 34 new Vpr substrates, together with further, extensive Vpr-dependent changes (downregulated and upregulated proteins) in HIV-infected CEM-T4s^39^. For comparison with this study, a list of 1,388 proteins concordantly regulated by Vpr (q<0.05) in the context of both viral infection and Vpr-bearing virus-like particles was compiled from published datasets. RUNX1 target genes were previously reported to be regulated by Vif at a transcriptional level because of competition for CBFβ binding^40^. For comparison with this study, a published list of 155 genes with RUNX1-associated regulatory domains exhibiting Vif-dependent differential gene expression in Jurkat T cells after 4 or 6 hrs of PMA and PHA treatment was used. Curated lists of ISGs have been previously described^90,91^. For comparison with this study, a list of 377 ISGs was compiled from these studies. A recent study quantitated 7,761 proteins in FACS-sorted T cells at a single time point 96 hrs post-infection with an R5-tropic, GFP-expressing Nef-deficient virus^70^. The comparator is GFP negative rather than mock-infected cells (equivalent to SBP-ΔLNGFR negative cells in this study), and the full dataset is not available. For comparison with this study, a published list of 1,551 differentially expressed proteins (q<0.05) was therefore used.

#### Flow cytometry

For primary T cells, CD3/CD28 Dynabeads were first removed using a DynaMag-2 magnet (Invitrogen). Typically 2e5 washed cells were incubated for 15 mins in 100 μL PBS with the indicated fluorochrome-conjugated antibody. All steps were performed on ice or at 4°C and stained cells were fixed in PBS/1% paraformaldehyde.

#### Immunoblotting

Cells were lysed in PBS/2% SDS supplemented with Halt Protease Inhibitor Cocktail (Thermo Scientific) and Halt Phosphatase Inhibitor Cocktail (Thermo Scientific) for 10 mins at RT. Benzonase (Sigma) was included to reduce lysate viscosity. Post-nuclear supernatants were heated in Laemelli Loading Buffer for 25 mins at 50°C, separated by SDS-PAGE and transferred to Immobilon-P membrane (Millipore). Membranes were blocked in PBS/5% non-fat dried milk (Marvel)/0.2% Tween and probed with the indicated primary antibody overnight at 4°C. Reactive bands were visualised using HRP-conjugated secondary antibodies and SuperSignal West Pico or Dura chemiluminescent substrates (Thermo Scientific). Typically 10-20μg total protein was loaded per lane (Pierce BCA Protein Assay kit).

#### Antibodies used in this study

| Antibody | Source* | Catalogue number | Application† |
| --- | --- | --- | --- |
| BV421-conjugated anti-CD4 | BioLegend | 317434 | FC |
| PE-conjugated anti-CD4 | BioLegend | 561843 | FC |
| PE-conjugated anti-tetherin | BioLegend | 348405 | FC |
| AF647-conjugated anti-LNGFR | BioLegend | 345114 | FC |
| FITC-conjugated anti-LNGFR | BioLegend | 345103 | FC |
| Anti-PPP2R5D | Abcam | ab188323 | IB |
| Anti-Vif | NIH ARP | #6459 | IB |
| Anti-p24 | Abcam | ab9071 | IB |
| Anti-Nef | NIH ARP | 3689 | IB |
| Anti-PTPN22 (D6D1H) | CST | 14693 | IB |
| Anti-ARID5A | GeneTex | GTX361940 | IB |
| Anti-FMRP (FMR1) | CST | 4317 | IB |
| Anti-DPH7 | Atlas Antibodies | R10023 | IB |
| Anti-tubulin | CST | 3873 | IB |
| Anti-β-actin | Sigma | A5316 | IB |
| Anti-total AURKB | CST | 3094 | IB |
| Anti-phospho-AURK | CST | 2914 | IB |

\*ARP, AIDS Reagent Program; CST, Cell Signalling Technology.
†FC, flow cytometry; IB, immunoblot.

## Acknowledgments

This work was supported by the MRC (CSF MR/P008801/1 to NJM), NHSBT (WPA15-02 to NJM), the Wellcome Trust (PRF 210688/Z/18/Z to PJL), the NIHR Cambridge BRC, and a Wellcome Trust Strategic Award to CIMR. The authors thank Dr Reiner Schulte and the CIMR Flow Cytometry Core Facility team, and members of the Lehner laboratory for critical discussion.

## Supplementary Figures

**Supplementary Figure 1:**
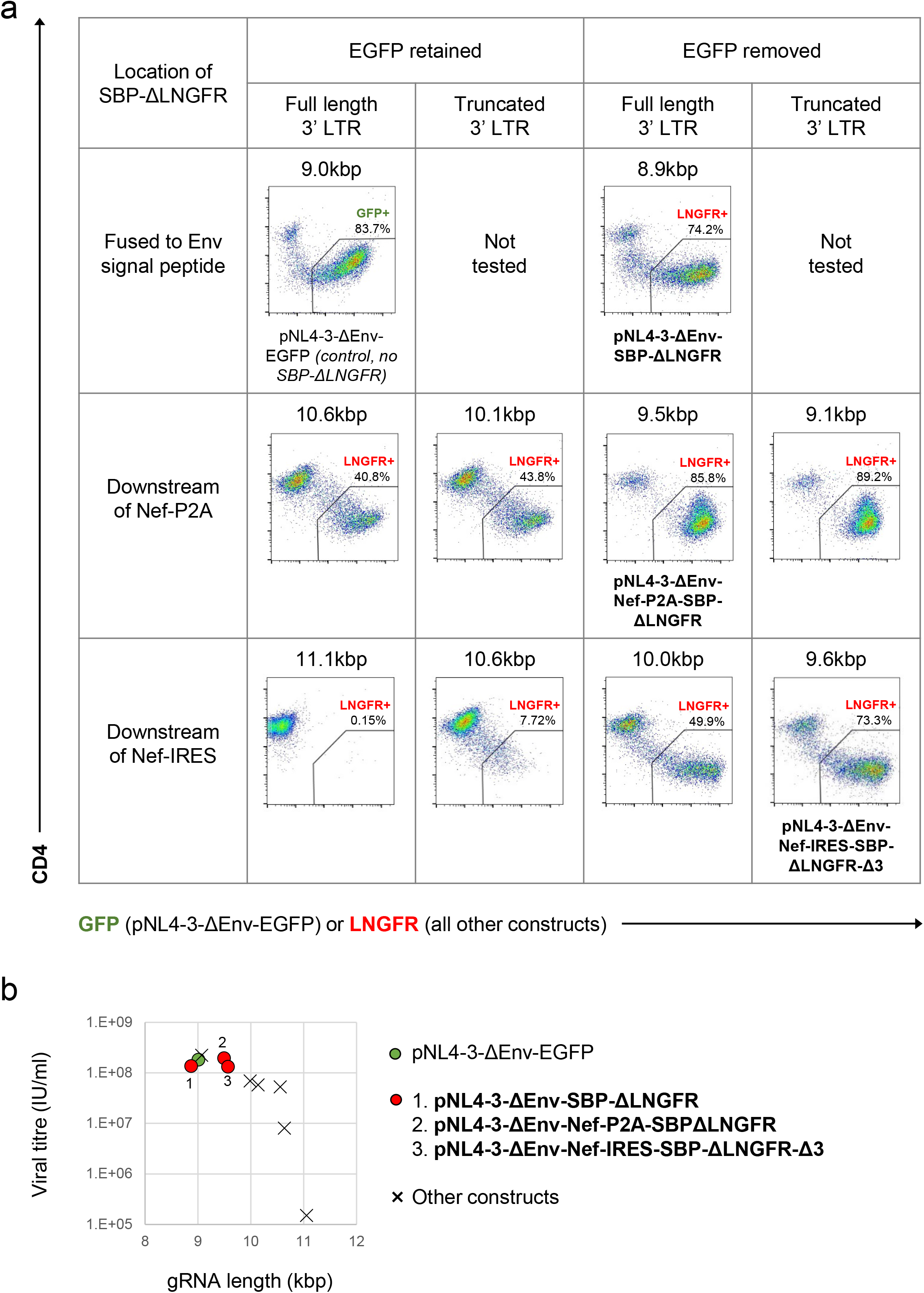
Initial screen of SBP-ΔLNGFR-expressing HIV viruses. **a, b**, Expression of GFP (pNL4-3-ΔEnv-EGFP only) or cell surface SBP-ΔLNGFR (all other constructs) and CD4 on CEM-T4s 48 hrs post-infection with indicated pNL4-3-ΔEnv-based viruses (**a**). Cells were stained with anti-LNGFR and anti-CD4 antibodies and analysed by flow cytometry. gRNA length is shown for each construct, and compared with functional viral titre derived from the % LNGFR+ cells (**b**). The 3 viruses selected for further testing are named/highlighted (bold text). pNL4-3-ΔEnv-EGFP was included as a control.

**Supplementary Figure 2:**
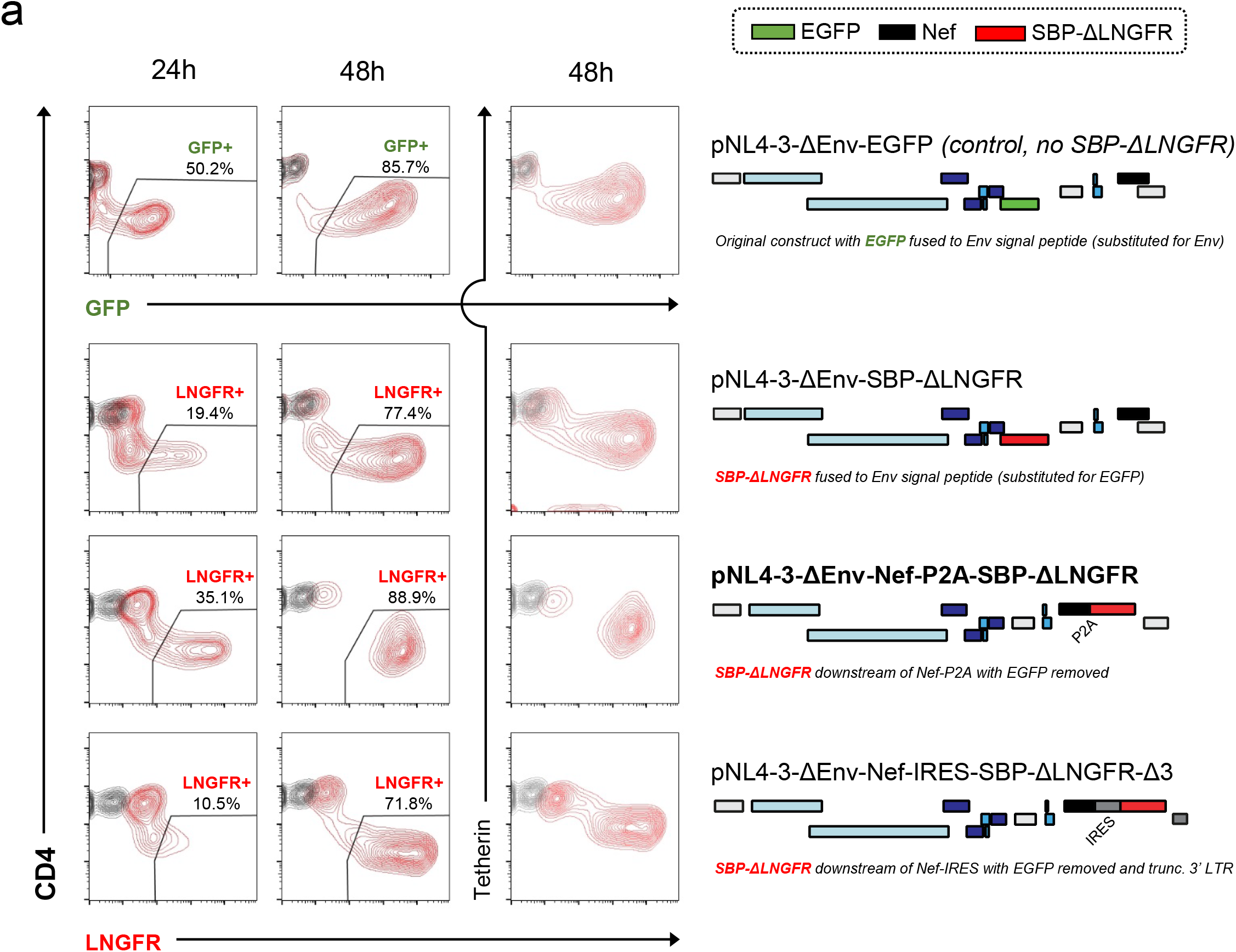
Time course evaluation of selected SBP-ΔLNGFR-expressing HIV viruses. **a**, Expression of GFP (pNL4-3-ΔEnv-EGFP only) or cell surface SBP-ΔLNGFR (all other constructs) and CD4 or tetherin in CEM-T4s 24 or 48 hrs post-infection with HIV-AFMACS. Cells were stained with anti-LNGFR and anti-CD4 or anti-tetherin antibodies at the indicated time points and analysed by flow cytometry. Schematics indicate the location/setting of SBP-ΔLNGFR within each provirus, with ORFs and non-coding features coloured as in **Fig. 1b** (but with Nef in black and EGFP in green). For simplicity, reading frames are drawn to match the HXB2 HIV-1 reference genome. The final HIV-AFMACS virus is highlighted (pNL4-3-ΔEnv-Nef-P2A-SBP-ΔLNGFR, bold text). pNL4-3-ΔEnv-EGFP was included as a control.

**Supplementary Figure 3:**
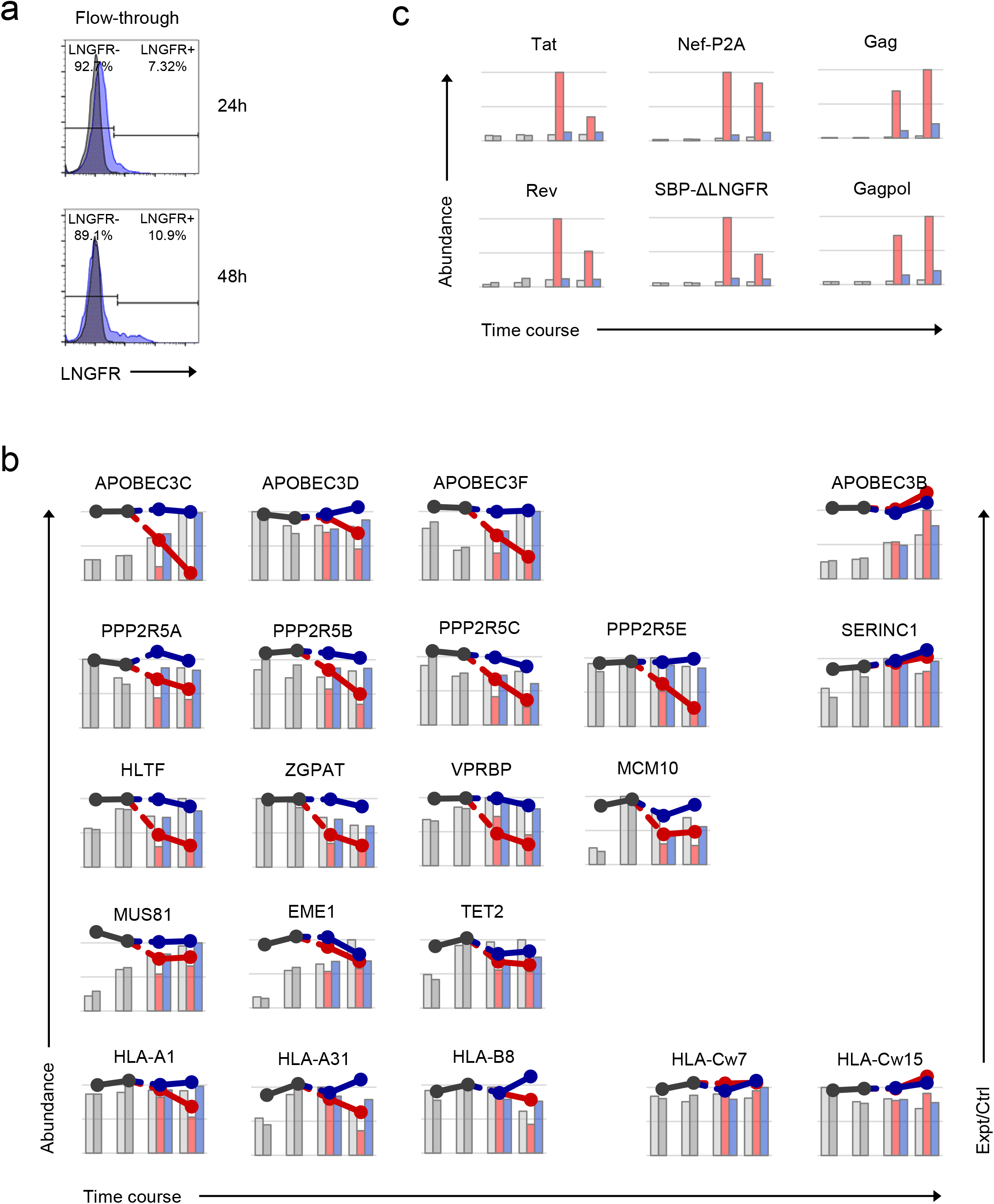
Additional controls for time course proteomic experiment. **a**, Uninfected (LNGFR-, flow-through) cells used for time course proteomic experiment (**Fig. 2a**). Corresponding HIV-infected (LNGFR+, selected) cells are shown in **Fig. 2b**. Cells were separated using AFMACS at the indicated time points post-infection with HIV-AFMACS, stained with anti-LNGFR and anti-CD4 antibodies and analysed by flow cytometry. **b, c**, Expression profiles of accessory protein targets (b; APOBEC3 and PPP2R5 family proteins, Vif; HLA-A/B alleles, Nef; other downregulated proteins, Vpr) and viral proteins (c) from time course proteomic experiment (**Fig. 2a**). Axes, scales and colours are as in **Fig. 2c** with the exception of APOBEC3C (range of log2(Expt/Ctrl) 0 to −4 rather than 0 to −2.5). SERINC1, APOBEC3B and HLA-C alleles are included as controls. Only HLA alleles quantitated by >1 unique peptide are shown. Since viral proteins are not expressed in control (Ctrl) cells, log2(ratio)s of abundances in Expt:Ctrl cells are not shown. Only canonical isoforms of PPP2R5C (Q13362) and ZGPAT (Q8N5A5) are shown.

**Supplementary Figure 4:**
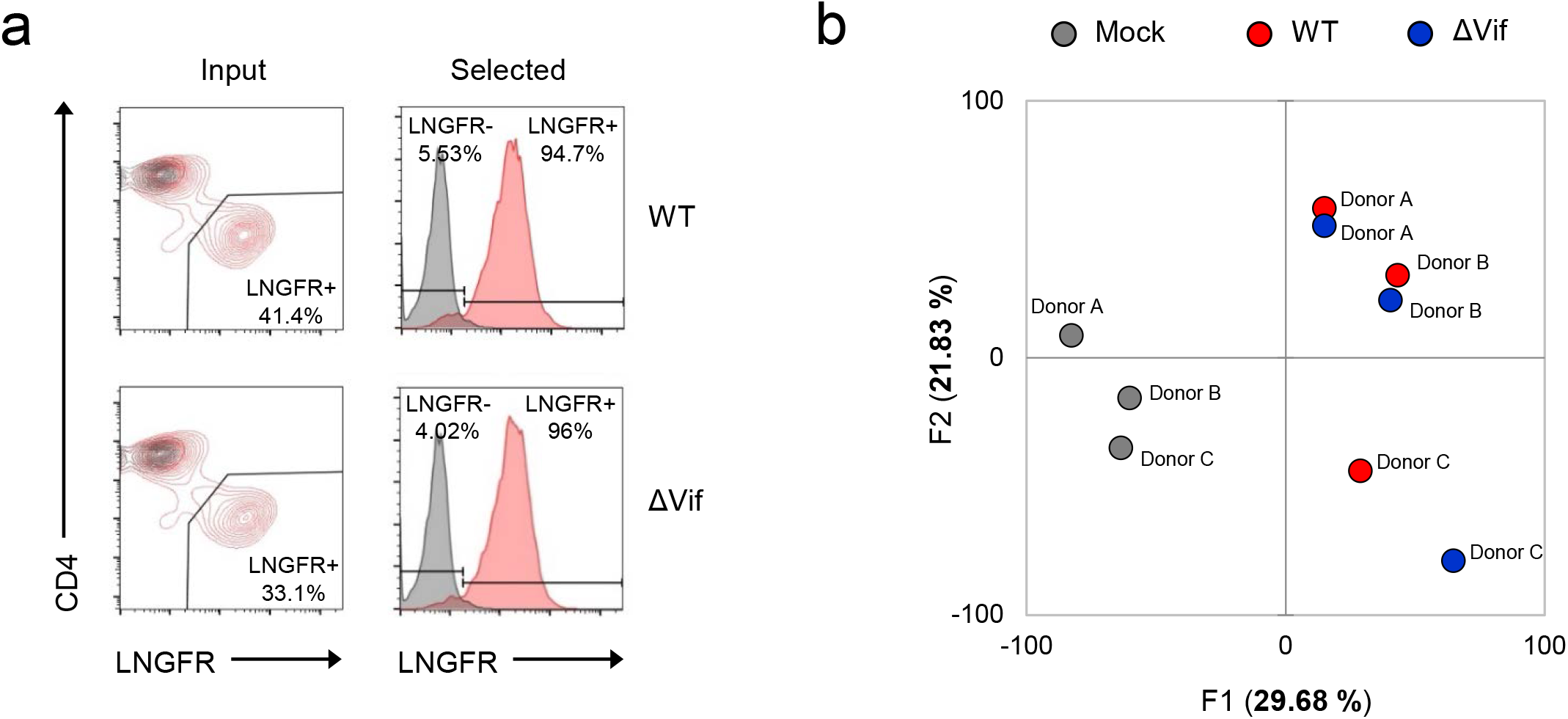
Additional controls for single time point proteomic experiment. **a**, AFMACS-based enrichment of HIV-infected (LNGFR+) cells used for single time point proteomic experiment (**Fig. 3a**). Cells were stained with anti-LNGFR and anti-CD4 antibodies and analysed by flow cytometry. Representative data is shown from donor B, with summary data in **Fig. 3b**. **b**, Principle component analysis of mock-infected (grey), WT (red) and ΔVif (blue) HIV-infected samples from single time point proteomic experiment (**Fig. 3a**). The correlation matrix was analysed for 8,795 cellular and viral proteins quantitated in cells from all 3 donors (no missing values). Visually-identical results were obtained with/without viral proteins.

**Supplementary Figure 5:**
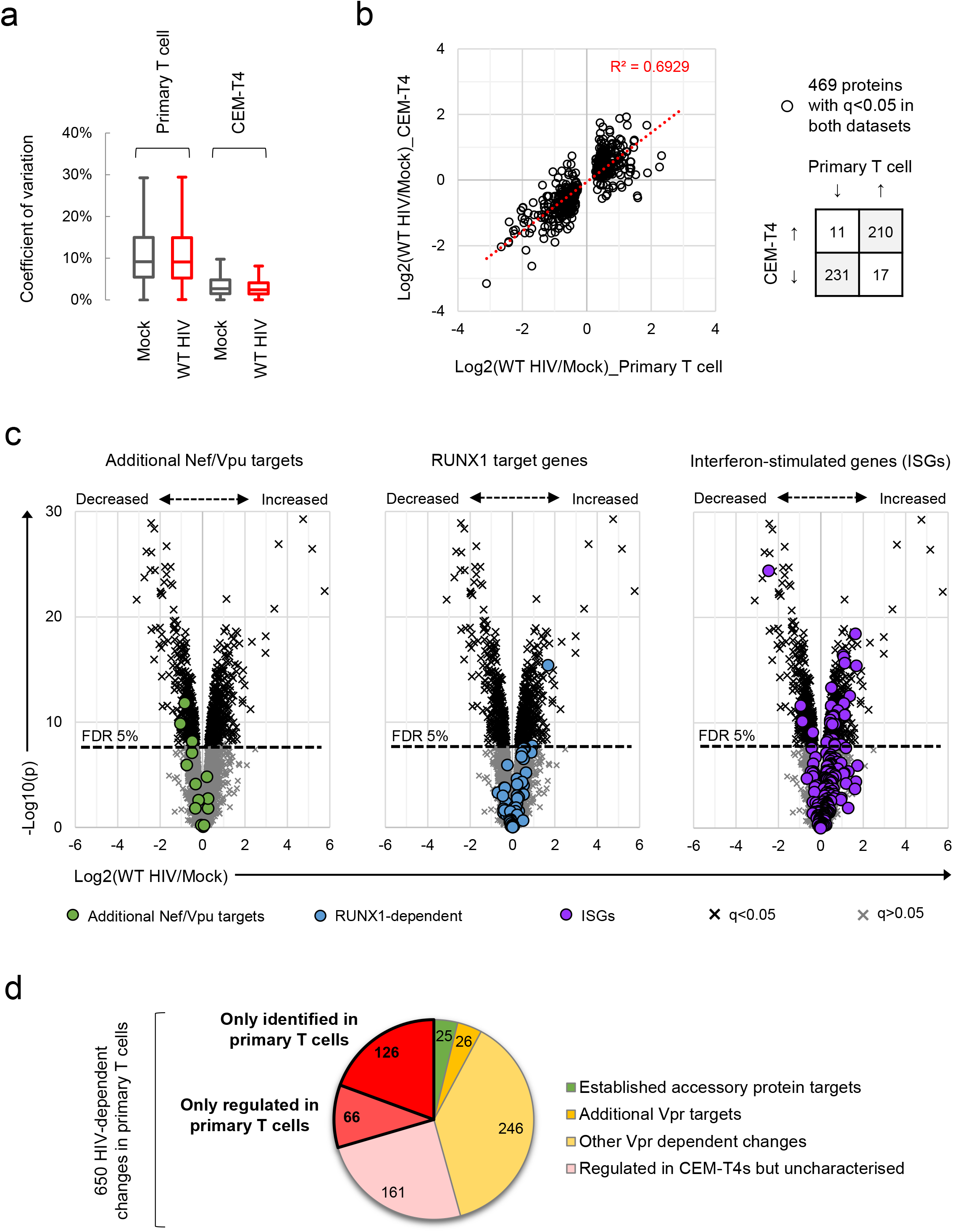
Comparisons of HIV-dependent changes in primary T cells with other datasets. **a**, Between-sample coefficients of variation (%) for all protein abundances from primary T cells (single time point proteomic experiment, **Fig. 3a**) or CEM-T4s^1^. Boxplots show median, interquartile range and Tukey whiskers for mock-infected (grey) and WT HIV-infected (red) cells. **b**, Protein abundances in WT HIV-infected vs mock-infected cells from primary T cells (x axis; single time point proteomic experiment, **Fig. 3a**) or CEM-T4s (y axis)^1^. Fold changes are compared for proteins with q<0.05 in both datasets. **c**, Protein abundances in WT HIV-infected vs mock-infected cells from single time point proteomic experiment (**Fig. 3a**), with details for volcano plots as in **Fig. 3c**. Proteins highlighted in each plot are summarised in the legend. Additional Nef/Vpu targets (green circles) comprise (alphabetical order, grouped by study): CCR7^2^, CD37/CD53/CD63/CD81/CD82^3,4^, CD99 (2 quantitated gene products, H7C2F2 and P14209)/PLP2/UBE2L6^5^, ICAM1/3^6^, NTB-A^7^, PVR^8,9^, SELL^10^. RUNX1 target genes (blue circles) were previously reported to be regulated by Vif at a transcriptional level because of competition for CBFβ binding^11^. ISGs (purple circles) were previously curated from published microarray data sets from IFN-treated cells^12,13^. **d**, Mechanisms underlying significant HIV-dependent proteins changes (q<0.05) in single time point proteomic experiment (**Fig. 3a**). Established accessory protein targets from **Fig. 3c** (left panel, 22 proteins) and c (left panel, 3 proteins) and additional Vpr substrates and Vpr-dependent changes from **Fig. 3c** (middle panel) are shown. Amongst the 32 additional Vpr substrates depleted in primary T cells, 26 had q<0.05. Remaining proteins are categorised based on the identification plus/minus HIV-dependent regulation in a previous, similar experiment using CEM-T4s^14^. Of 131 proteins not identified in the comparator CEM-T4 dataset, one is known Vpr target SMUG1^15^ and 4 other proteins were found to be regulated by Vpr in other experiments using CEM-T4s (ZNF512B, ATXN7, CLUH and SLC39A3)^14^. Further details on comparator datasets used in this figure are provided in the **Methods**.

**Supplementary Figure 6:**
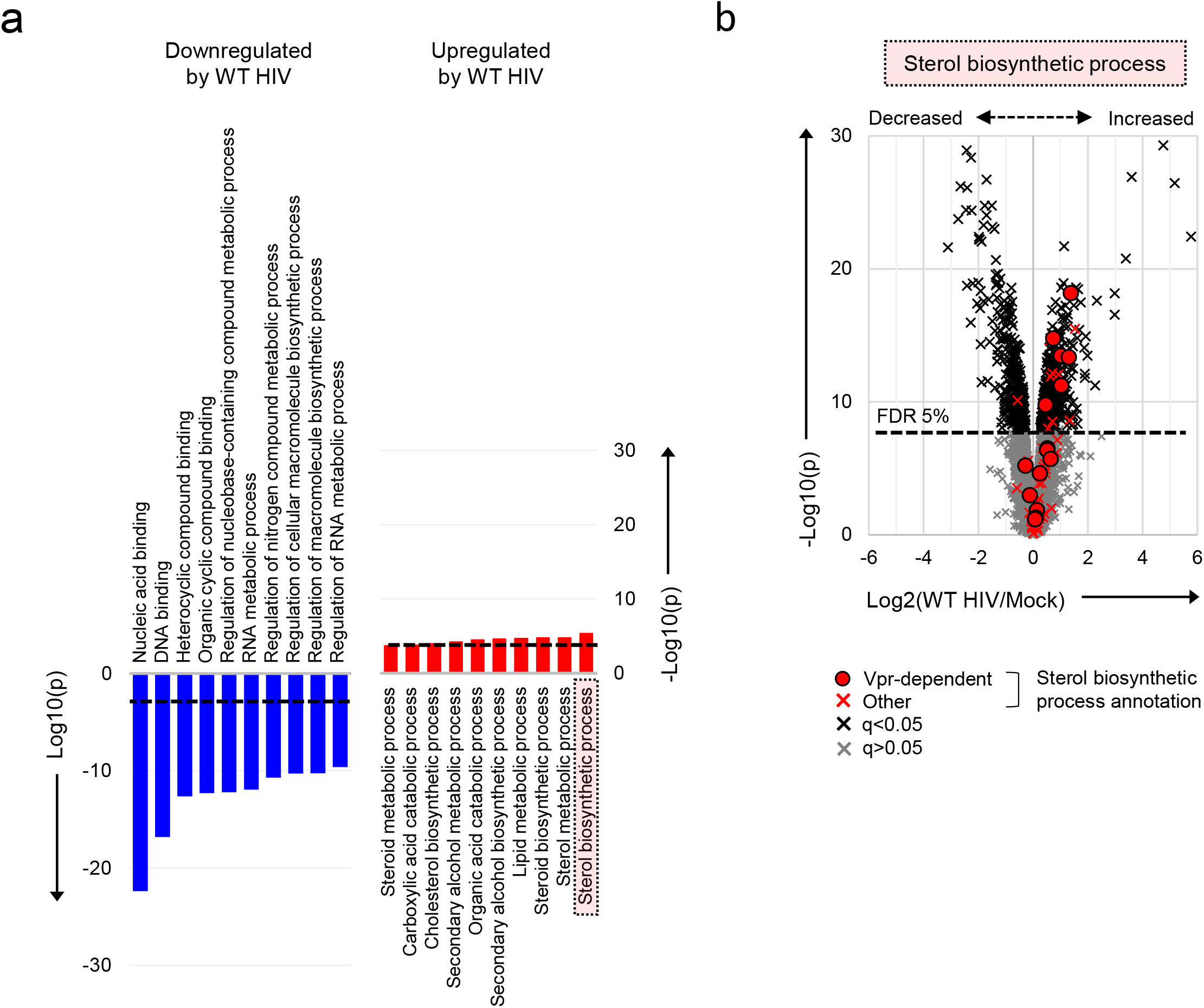
Pathways regulated by HIV in primary T cells. **a**, Gene Ontology (GO) functional annotations enriched amongst downregulated (blue) or upregulated (red) proteins with q<0.05 in WT HIV-infected vs mock-infected cells from single time point proteomic experiment (**Fig. 3a**). The 10 most enriched terms (ranked by p value) are shown in each case, with a Benjamini-Hochberg FDR threshold of 5% indicated (dashed line). **b**, Protein abundances in WT HIV-infected vs mock-infected cells from single time point proteomic experiment (**Fig. 3a**), with details for volcano plot as in **Fig. 3c**. 57 proteins annotated with the GO term “sterol biosynthetic process” (GO:0016126) are highlighted in red. Amongst these, 15 proteins are regulated by Vpr in CEM-T4s (circles)^14^.

**Supplementary Figure 7:**
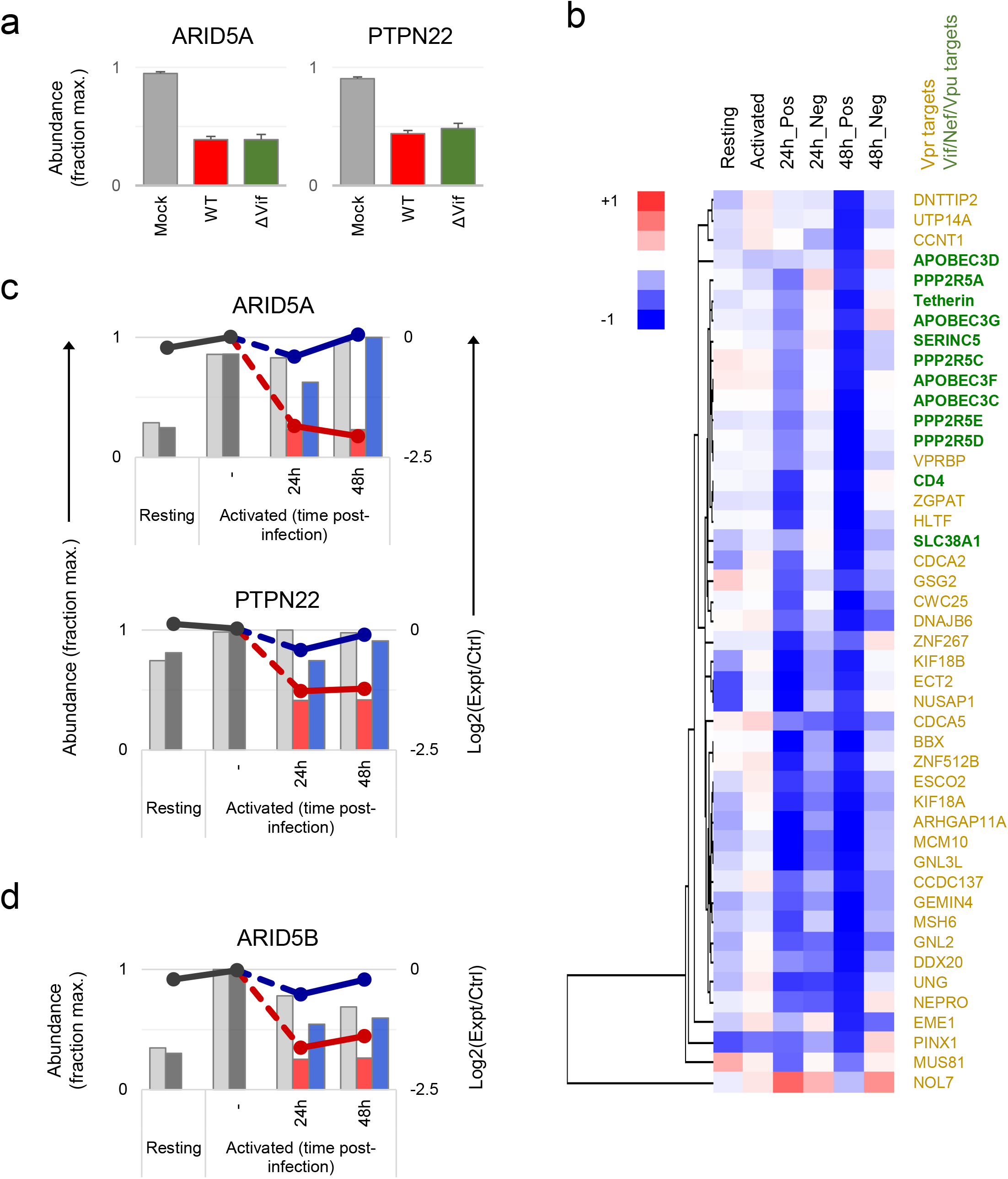
Temporal clustering of HIV accessory protein targets. **a**, Abundances of ARID5A and PTPN22 in mock-infected (grey), WT HIV-infected (red) and ΔVif HIV-infected (green) primary T cells from single time point proteomic experiment (**Fig. 3a**). Relative abundances (fraction of maximum) with 95% confidence intervals are shown. **b**, Hierarchical cluster analysis of 45 accessory protein targets according to profiles of temporal expression from time course proteomic experiment (**Fig. 2a**). Vpr (gold) vs other accessory protein (Vif/Nef/Vpu; green) targets are highlighted. The heatmap shows range-scaled log2(ratio)s of abundances in experimental (Expt):control (Ctrl) cells for each condition/time point. Unscaled data for the same proteins are shown in **Fig. 2e**. **c, d**, Expression profiles of ARID5A, PTPN22 and AIRD5B from time course proteomic experiment (**Fig. 2a**). Axes, scales and colours are as in **Fig. 2c**.

**Supplementary Figure 8:**
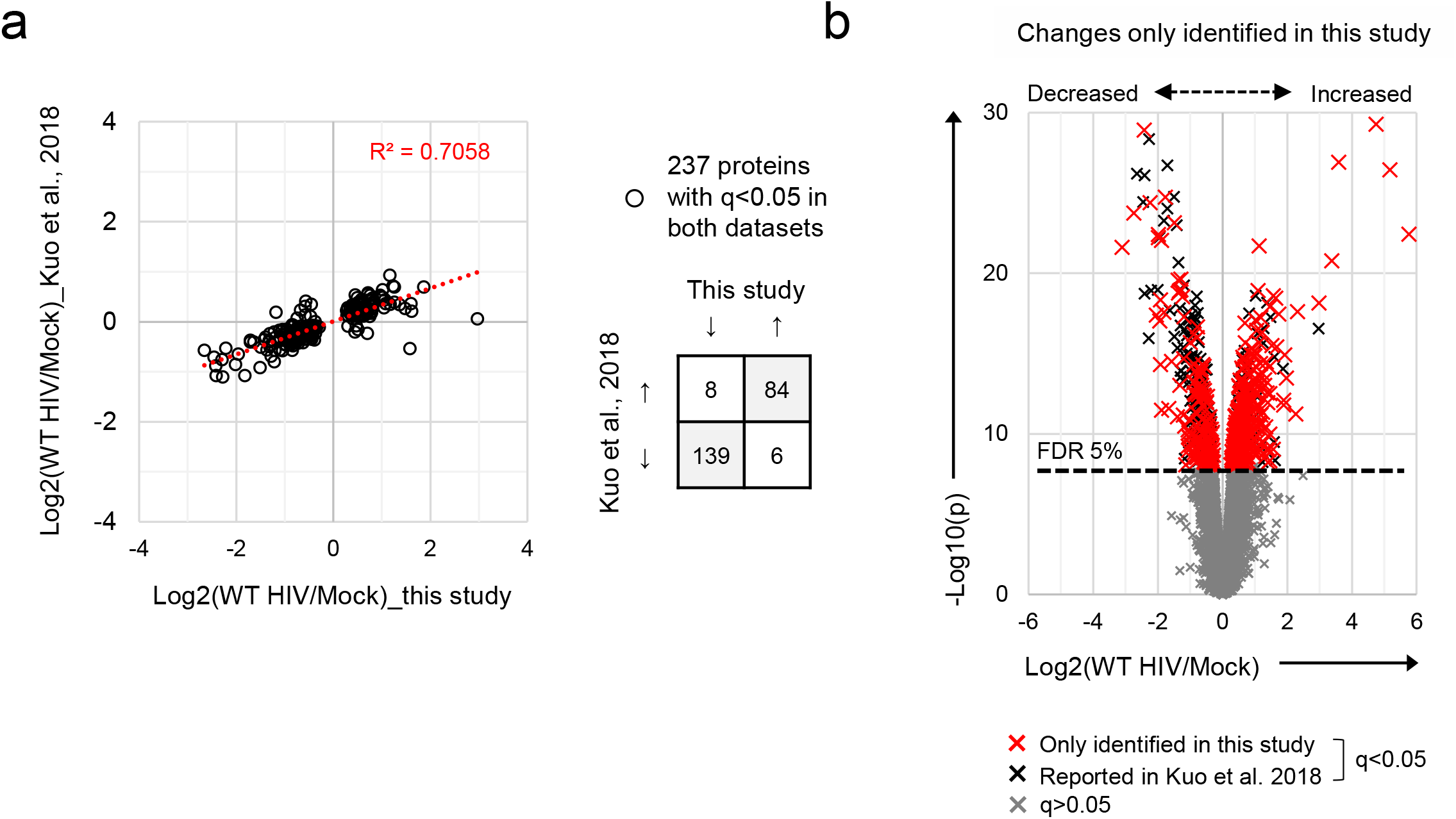
Comparison of this study with Kuo et al., 2018^16^. **a**, Protein abundances in WT HIV-infected vs mock-infected cells from this study (x axis; single time point proteomic experiment, **Fig. 3a**) or Kuo et al., 2018^16^ (y axis). Fold changes are compared for proteins with q<0.05 in both datasets. **b**, Protein abundances in WT HIV-infected vs mock-infected cells from single time point proteomic experiment (**Fig. 3a**), with details for volcano plot as in **Fig. 3c**. Proteins with q<0.05 not reported by Kuo et al., 2018^16^ are highlighted in red.

### Supplementary Tables

**Supplementary Table 1:**
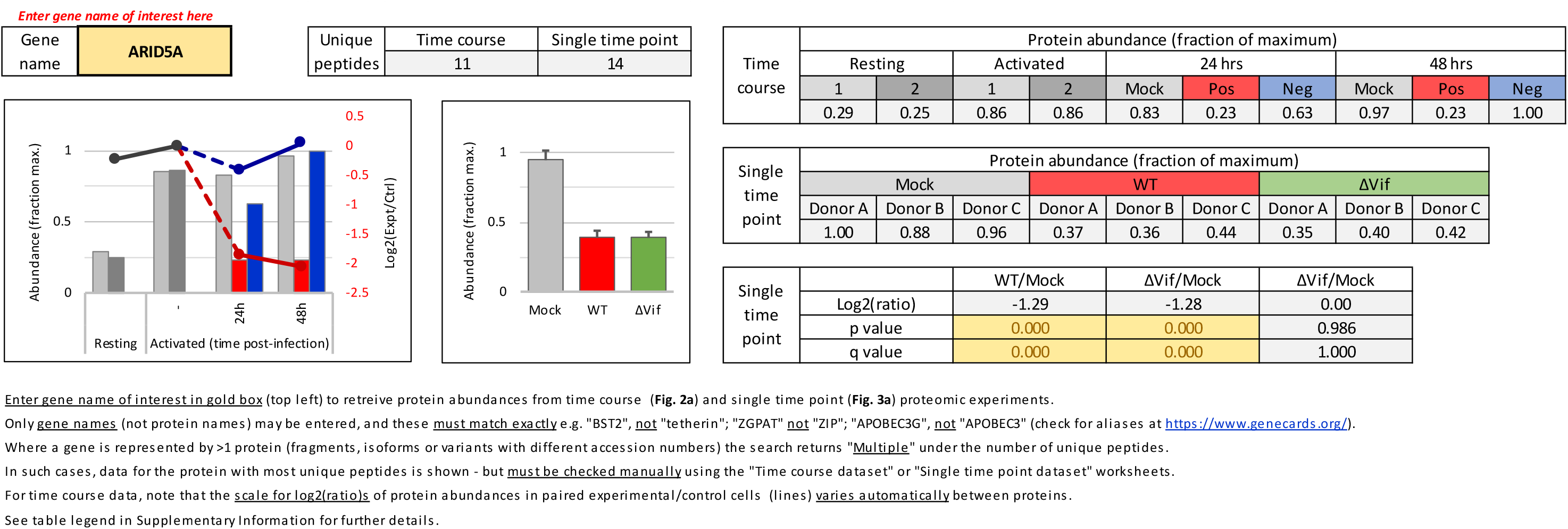
Functional proteomic atlas of HIV-infection in primary human CD4+ T cells. Interactive spreadsheet enabling generation of temporal profiles of protein abundance for any quantitated genes of interest (“Gene search and plots” worksheet). Time course data (cells from **Fig. 2a**) are presented as in **Fig. 2c**, with relative protein abundances (fraction of maximum) for each condition depicted by bars, and log2(ratio)s of protein abundances in paired experimental/control cells from each condition/time point depicted by lines (grey, resting/activated; red, LNGFR+, infected; blue, LNGFR-, uninfected). Single time point data (cells from **Fig. 3a**) are presented as in **Fig. 4b**, with relative protein abundances (fraction of maximum, mean plus 95% CIs) for each condition depicted by bars (grey, mock; red, WT HIV; green, ΔVif HIV). The number of unique peptides is shown for each protein/experiment, with most confidence reserved for proteins with values >1. For the single time point experiment, p values (unadjusted) and q values (Benjamini-Hochberg FDR-adjusted) are shown (highlighted in gold if <0.05). The complete (unfiltered) proteomic datasets (“Time course dataset” and “Single time point dataset” worksheets) are also included.

### Supplementary Methods

#### gBlock sequences

##### gBlock #1

~~~
tgtttatccatttcagaattgggtgtcgacatagcagaataggcgttactcgacagaggagagcaagaaatggagccagtaga
tcctagactagagccctggaagcatccaggaagtcagcctaaaactgcttgtaccaattgctattgtaaaaagtgttgctttc
attgccaagtttgtttcatgacaaaagccttaggcatctcctatggcaggaagaagcggagacagcgacgaagagctcatcag
aacagtcagactcatcaagcttctctatcaaagcagtaagtagtacatgtaatgcaacctataatagtagcaatagtagcatt
agtagtagcaataataatagcaatagttgtgtggtccatagtaatcatagaatataggaaaatattaagacaaagaaaaatag
acaggttaattgatagactaatagaaagagcagaagacagtggcaatgagagtgaaggagaagtatcagcacttgtggagatg
ggggtggaaatggggcaccatgctccttgggatattgatgatctgtagtgctacagaaaaattgtgggtcacagtctattatg
gggtacctgtcgcggccgcagggtcagggatggacgaaaagacaaccggatggcgaggaggacacgtggtcgagggactggca
ggagagctggaacagctgcgggctagactggaacaccatcctcagggacagcgagagcccggaagtggaaaggaagcctgccc
cacagggctgtacactcattctggagaatgctgtaaagcttgtaacctgggagagggagtggcacagccatgcggagccaatc
agactgtgtgcgagccttgtctggactccgtcacattctctgatgtggtcagtgccacagaaccttgcaagccatgtactgag
tgcgtgggcctgcagtctatgagtgctccttgtgtggaggctgacgatgcagtctgccggtgtgcatacggatactatcagga
cgagactaccggcagatgtgaagcttgcagggtgtgtgaggcaggctcagggctggtctttagctgccaggataaacagaaca
ccgtgtgcgaggaatgtcctgacgggacatatagcgatgaggccaatcacgtggacccctgcctgccttgtactgtgtgcgag
gataccgaaaggcagctgcgcgaatgtaccagatgggcagacgccgagtgcgaggaaatcccagggcgatggattactcggtc
caccccccctgaaggatcagacagcaccgcaccatctacacaggagccagaagcaccaccagagcaggatctgatcgcctcca
ccgtggctggcgtggtcacaactgtcatggggagctctcagccagtggtcacccggggcaccacagataacctgattcccgtg
tattgctccatcctggcagccgtcgtcgtcggactggtggcatacatcgccttcaagcggtgagctagccgtaacgatgttca
tcaaatattactgggctgctat
~~~

##### gBlock #2

~~~
tcaggaactaaagaatagtgctgttaacttgctcaatgccacagccatagcagtagctgaggggacagatagggttatagaag
tattacaagcagcttatagagctattcgccacatacctagaagaataagacagggcttggaaaggattttgctataagatggg
ggggaaatggtctaagtcttccgtgataggctggcccgctgttcgagaaaggatgcgccgcgccgaaccagcggccgatggcg
taggtgccgtatctagggaccttgagaagcacggtgccataacgtcttctaacaccgcagccaacaatgcggcatgtgcgtgg
ttggaggcacaggaagaggaggaggtaggattcccggtaacgccacaagtacccttgcgccctatgacctacaaggccgccgt
ggacctgagccatttcctgaaggagaaaggtggcttggaagggcttatccactcccaaagacgccaggatattcttgaccttt
ggatctaccacacacagggttactttcccgactggcaaaattatacgcccggtcccggtgtacggtatcctcttacttttggg
tggtgctataaactcgtgccggtggagccagataaggtagaggaggccaataaaggtgagaacacaagtttgcttcaccccgt
ttcccttcatggtatggacgacccagaacgggaagttcttgaatggcggttcgatagcaggttggcatttcatcatgtcgcac
gggagcttcaccccgagtactttaaaaattgtgcggccgcaggaagcggagctactaacttcagcctgctgaagcaggctgga
gacgtggaggagaaccctggacctatgggctggtcatgtatcattctgtttctggtcgcaaccgcaactggagtgcattcaca
ggtgcagctcagcgctgggtcagggatggacgaaaagacaaccggatggcgaggaggacacgtggtcgagggactggcaggag
agctggaacagctgcgggctagactggaacaccatcctcagggacagcgagagcccggaagtggagcgatcgcgaaggaagcc
tgccccacagggctgtacactcattctggagaatgctgtaaagcttgtaacctgggagagggagtggcacagccatgcggagc
caatcagactgtgtgcgagccttgtctggactccgtcacattctctgatgtggtcagtgccacagaaccttgcaagccatgta
ctgagtgcgtgggcctgcagtctatgagtgctccttgtgtggaggctgacgatgcagtctgccggtgtgcatacggatactat
caggacgagactaccggcagatgtgaagcttgcagggtgtgtgaggcaggctcagggctggtctttagctgccaggataaaca
gaacaccgtgtgcgaggaatgtcctgacgggacatatagcgatgaggccaatcacgtggacccctgcctgccttgtactgtgt
gcgaggataccgaaaggcagctgcgcgaatgtaccagatgggcagacgccgagtgcgaggaaatcccagggcgatggattact
cggtccaccccccctgaaggatcagacagcaccgcaccatctacacaggagccagaagcaccaccagagcaggatctgatcgc
ctccaccgtggctggcgtggtcacaactgtcatggggagctctcagccagtggtcacccggggcaccacagataacctgattc
ccgtgtattgctccatcctggcagccgtcgtcgtcggactggtggcatacatcgccttcaagcggtgactcgagacctagaaa
aacatggagcaatca
~~~

##### gBlock #3

~~~
ggagtcaggaactaaagaatagtgctgttaacttgctcaatgccacagccatagcagtagctgaggggacagatagggttata
gaagtattacaagcagcttatagagctattcgccacatacctagaagaataagacagggcttggaaaggattttgctataaga
tgggggggaaatggtctaagtcttccgtgataggctggcccgctgttcgagaaaggatgcgccgcgccgaaccagcggccgat
ggcgtaggtgccgtatctagggaccttgagaagcacggtgccataacgtcttctaacaccgcagccaacaatgcggcatgtgc
gtggttggaggcacaggaagaggaggaggtaggattcccggtaacgccacaagtacccttgcgccctatgacctacaaggccg
ccgtggacctgagccatttcctgaaggagaaaggtggcttggaagggcttatccactcccaaagacgccaggatattcttgac
ctttggatctaccacacacagggttactttcccgactggcaaaattatacgcccggtcccggtgtacggtatcctcttacttt
tgggtggtgctataaactcgtgccggtggagccagataaggtagaggaggccaataaaggtgagaacacaagtttgcttcacc
ccgtttcccttcatggtatggacgacccagaacgggaagttcttgaatggcggttcgatagcaggttggcatttcatcatgtc
gcacgggagcttcaccccgagtactttaaaaattgttgatctagacgcccccccctaacgttactggccgaagccgcttggaa
taaggccggtgtgcgtttgtctatatgttattttccaccatattgccgtcttttggcaatgtgagggcccggaaacctggccc
tgtcttcttgacgagcattcctaggggtctttcccctctcgccaaaggaatgcaaggtctgttgaatgtcgtgaaggaagcag
ttcctctggaagcttcttgaagacaaacaacgtctgtagcgaccctttgcaggcagcggaaccccccacctggcgacaggtgc
ctctgcggccaaaagccacgtgtataagatacacctgcaaaggcggcacaaccccagtgccacgttgtgagttggatagttgt
ggaaagagtcaaatggctctcctcaagcgtattcaacaaggggctgaaggatgcccagaaggtaccccattgtatgggatctg
atctggggcctcggtgcacatgctttacatgtgtttagtcgaggttaaaaaacgtctaggccccccgaaccacggggacgtgg
ttttcctttgaaaaacacgatgataatatgggctggtcatgtatcattctgtttctggtcgcaaccgcaactggagtgcattc
acaggtgcagctcagcgctgggtcagggatggacgaaaagacaaccggatggcgaggaggacacgtggtcgagggactggcag
gagagctggaacagctgcgggctagactggaacaccatcctcagggacagcgagagcccggaagtggagcgatcgcgaaggaa
gcctgccccacagggctgtacactcattctggagaatgctgtaaagcttgtaacctgggagagggagtggcacagccatgcgg
agccaatcagactgtgtgcgagccttgtctggactccgtcacattctctgatgtggtcagtgccacagaaccttgcaagccat
gtactgagtgcgtgggcctgcagtctatgagtgctccttgtgtggaggctgacgatgcagtctgccggtgtgcatacggatac
tatcaggacgagactaccggcagatgtgaagcttgcagggtgtgtgaggcaggctcagggctggtctttagctgccaggataa
acagaacaccgtgtgcgaggaatgtcctgacgggacatatagcgatgaggccaatcacgtggacccctgcctgccttgtactg
tgtgcgaggataccgaaaggcagctgcgcgaatgtaccagatgggcagacgccgagtgcgaggaaatcccagggcgatggatt
actcggtccaccccccctgaaggatcagacagcaccgcaccatctacacaggagccagaagcaccaccagagcaggatctgat
cgcctccaccgtggctggcgtggtcacaactgtcatggggagctctcagccagtggtcacccggggcaccacagataacctga
ttcccgtgtattgctccatcctggcagccgtcgtcgtcggactggtggcatacatcgccttcaagcggtgactcgagacctag
aaaaacatggagcaatcacaag
~~~

##### gBlock #4

~~~
tgtttatccatttcagaattgggtgtcgacatagcagaataggcgttactcgacagaggagagcaagaaatggagccagtaga
tcctagactagagccctggaagcatccaggaagtcagcctaaaactgcttgtaccaattgctattgtaaaaagtgttgctttc
attgccaagtttgtttcatgacaaaagccttaggcatctcctatggcaggaagaagcggagacagcgacgaagagctcatcag
aacagtcagactcatcaagcttctctatcaaagcagtaagtagtacatgtaatgcaacctataatagtagcaatagtagcatt
agtagtagcaataataatagcaatagttgtgtggtccatagtaatcatagaatataggaaaatattaagacaaagaaaaatag
acaggttaattgatagactaatagaaagagcagaagacagtggcaacgagagtgaaggagaagtatcagcacttgtggagatg
ggggtggaaatggggcaccacgctccttgggatattgacgatctgtaggctagccgtaacgatgttcatcaaatattactggg
ctgctat
~~~

##### gBlock #5

~~~
atacatcgccttcaagcggtgactcgagtagccactttttaaaagaaaaggggggactggaagggctaattcactcccaaaga
agacaagatatcccatccggagtacttcaagaactgctgacatcgagcttgctacaagggactttccgctggggactttccag
ggaggcgtggcctgggcgggactggggagtggcgagccctcagatgctgcatataagcagctgctttttgcctgtactgggtc
tctctggttagaccagatctgagcctgggagctctctggctaactagggaacccactgcttaagcctcaataaagcttgcctt
gagtgcttcaagtagtgtgtgcccgtctgttgtgtgactctggtaactagagatccctcagacccttttagtcagtgtggaaa
atctctagcacccaggaggtagaggttgcagtgagccaagatcgcgccactgcattccagcctgggcaagaaaacaagactgt
ctaaaataataataataagttaagggtattaaatatatttatacatggaggtcataaaaatatatatatttgggctgggcgca
gtggctcacacctgcgcccggccctttgggaggccgaggcaggtggatcacctgagtttgggagttccagaccagcctgacca
acatggagaaaccccttctctgtgtatttttagtagattttattttatgtgtattttattcacaggtatttctggaaaactga
aactgtttttcctctactctgataccacaagaatcatcagcacagaggaagacttctgtgatcaaatgtggtgggagagggag
gttttcaccagcacatgagcagtcagttctgccgcagactcggcgggtgtccttcggttcagttccaacaccgcctgcctgga
gagaggtcagaccacagggtgagggctcagtccccaagacataaacacccaagacataaacaccccaggtccaccccgcctgc
tgcccaggcagagccgattcaccaagacgggaattaggatagagaaagagtaagtcacacagagccggctgtgcgggagaacg
gagttcta
~~~

##### HIV-AFMACS (pNL4-3-ΔEnv-Nef-P2A-SBP-ΔLNGFR) complete sequence

~~~
tggaagggctaatttggtcccaaaaaagacaagagatccttgatctgtggatctaccacacacaaggctacttccctgattgg
cagaactacacaccagggccagggatcagatatccactgacctttggatggtgcttcaagttagtaccagttgaaccagagca
agtagaagaggccaatgaaggagagaacaacagcttgttacaccctatgagccagcatgggatggaggacccggagggagaag
tattagtgtggaagtttgacagcctcctagcatttcgtcacatggcccgagagctgcatccggagtactacaaagactgctga
catcgagctttctacaagggactttccgctggggactttccagggaggtgtggcctgggcgggactggggagtggcgagccct
cagatgctacatataagcagctgctttttgcctgtactgggtctctctggttagaccagatctgagcctgggagctctctggc
taactagggaacccactgcttaagcctcaataaagcttgccttgagtgctcaaagtagtgtgtgcccgtctgttgtgtgactc
tggtaactagagatccctcagacccttttagtcagtgtggaaaatctctagcagtggcgcccgaacagggacttgaaagcgaa
agtaaagccagaggagatctctcgacgcaggactcggcttgctgaagcgcgcacggcaagaggcgaggggcggcgactggtga
gtacgccaaaaattttgactagcggaggctagaaggagagagatgggtgcgagagcgtcggtattaagcgggggagaattaga
taaatgggaaaaaattcggttaaggccagggggaaagaaacaatataaactaaaacatatagtatgggcaagcagggagctag
aacgattcgcagttaatcctggccttttagagacatcagaaggctgtagacaaatactgggacagctacaaccatcccttcag
acaggatcagaagaacttagatcattatataatacaatagcagtcctctattgtgtgcatcaaaggatagatgtaaaagacac
caaggaagccttagataagatagaggaagagcaaaacaaaagtaagaaaaaggcacagcaagcagcagctgacacaggaaaca
acagccaggtcagccaaaattaccctatagtgcagaacctccaggggcaaatggtacatcaggccatatcacctagaacttta
aatgcatgggtaaaagtagtagaagagaaggctttcagcccagaagtaatacccatgttttcagcattatcagaaggagccac
cccacaagatttaaataccatgctaaacacagtggggggacatcaagcagccatgcaaatgttaaaagagaccatcaatgagg
aagctgcagaatgggatagattgcatccagtgcatgcagggcctattgcaccaggccagatgagagaaccaaggggaagtgac
atagcaggaactactagtacccttcaggaacaaataggatggatgacacataatccacctatcccagtaggagaaatctataa
aagatggataatcctgggattaaataaaatagtaagaatgtatagccctaccagcattctggacataagacaaggaccaaagg
aaccctttagagactatgtagaccgattctataaaactctaagagccgagcaagcttcacaagaggtaaaaaattggatgaca
gaaaccttgttggtccaaaatgcgaacccagattgtaagactattttaaaagcattgggaccaggagcgacactagaagaaat
gatgacagcatgtcagggagtggggggacccggccataaagcaagagttttggctgaagcaatgagccaagtaacaaatccag
ctaccataatgatacagaaaggcaattttaggaaccaaagaaagactgttaagtgtttcaattgtggcaaagaagggcacata
gccaaaaattgcagggcccctaggaaaaagggctgttggaaatgtggaaaggaaggacaccaaatgaaagattgtactgagag
acaggctaattttttagggaagatctggccttcccacaagggaaggccagggaattttcttcagagcagaccagagccaacag
ccccaccagaagagagcttcaggtttggggaagagacaacaactccctctcagaagcaggagccgatagacaaggaactgtat
cctttagcttccctcagatcactctttggcagcgacccctcgtcacaataaagataggggggcaattaaaggaagctctatta
gatacaggagcagatgatacagtattagaagaaatgaatttgccaggaagatggaaaccaaaaatgatagggggaattggagg
ttttatcaaagtaagacagtatgatcagatactcatagaaatctgcggacataaagctataggtacagtattagtaggaccta
cacctgtcaacataattggaagaaatctgttgactcagattggctgcactttaaattttcccattagtcctattgagactgta
ccagtaaaattaaagccaggaatggatggcccaaaagttaaacaatggccattgacagaagaaaaaataaaagcattagtaga
aatttgtacagaaatggaaaaggaaggaaaaatttcaaaaattgggcctgaaaatccatacaatactccagtatttgccataa
agaaaaaagacagtactaaatggagaaaattagtagatttcagagaacttaataagagaactcaagatttctgggaagttcaa
ttaggaataccacatcctgcagggttaaaacagaaaaaatcagtaacagtactggatgtgggcgatgcatatttttcagttcc
cttagataaagacttcaggaagtatactgcatttaccatacctagtataaacaatgagacaccagggattagatatcagtaca
atgtgcttccacagggatggaaaggatcaccagcaatattccagtgtagcatgacaaaaatcttagagccttttagaaaacaa
aatccagacatagtcatctatcaatacatggatgatttgtatgtaggatctgacttagaaatagggcagcatagaacaaaaat
agaggaactgagacaacatctgttgaggtggggatttaccacaccagacaaaaaacatcagaaagaacctccattcctttgga
tgggttatgaactccatcctgataaatggacagtacagcctatagtgctgccagaaaaggacagctggactgtcaatgacata
cagaaattagtgggaaaattgaattgggcaagtcagatttatgcagggattaaagtaaggcaattatgtaaacttcttagggg
aaccaaagcactaacagaagtagtaccactaacagaagaagcagagctagaactggcagaaaacagggagattctaaaagaac
cggtacatggagtgtattatgacccatcaaaagacttaatagcagaaatacagaagcaggggcaaggccaatggacatatcaa
atttatcaagagccatttaaaaatctgaaaacaggaaagtatgcaagaatgaagggtgcccacactaatgatgtgaaacaatt
aacagaggcagtacaaaaaatagccacagaaagcatagtaatatggggaaagactcctaaatttaaattacccatacaaaagg
aaacatgggaagcatggtggacagagtattggcaagccacctggattcctgagtgggagtttgtcaatacccctcccttagtg
aagttatggtaccagttagagaaagaacccataataggagcagaaactttctatgtagatggggcagccaatagggaaactaa
attaggaaaagcaggatatgtaactgacagaggaagacaaaaagttgtccccctaacggacacaacaaatcagaagactgagt
tacaagcaattcatctagctttgcaggattcgggattagaagtaaacatagtgacagactcacaatatgcattgggaatcatt
caagcacaaccagataagagtgaatcagagttagtcagtcaaataatagagcagttaataaaaaaggaaaaagtctacctggc
atgggtaccagcacacaaaggaattggaggaaatgaacaagtagataaattggtcagtgctggaatcaggaaagtactatttt
tagatggaatagataaggcccaagaagaacatgagaaatatcacagtaattggagagcaatggctagtgattttaacctacca
cctgtagtagcaaaagaaatagtagccagctgtgataaatgtcagctaaaaggggaagccatgcatggacaagtagactgtag
cccaggaatatggcagctagattgtacacatttagaaggaaaagttatcttggtagcagttcatgtagccagtggatatatag
aagcagaagtaattccagcagagacagggcaagaaacagcatacttcctcttaaaattagcaggaagatggccagtaaaaaca
gtacatacagacaatggcagcaatttcaccagtactacagttaaggccgcctgttggtgggcggggatcaagcaggaatttgg
cattccctacaatccccaaagtcaaggagtaatagaatctatgaataaagaattaaagaaaattataggacaggtaagagatc
aggctgaacatcttaagacagcagtacaaatggcagtattcatccacaattttaaaagaaaaggggggattggggggtacagt
gcaggggaaagaatagtagacataatagcaacagacatacaaactaaagaattacaaaaacaaattacaaaaattcaaaattt
tcgggtttattacagggacagcagagatccagtttggaaaggaccagcaaagctcctctggaaaggtgaaggggcagtagtaa
tacaagataatagtgacataaaagtagtgccaagaagaaaagcaaagatcatcagggattatggaaaacagatggcaggtgat
gattgtgtggcaagtagacaggatgaggattaacacatggaaaagattagtaaaacaccatatgtatatttcaaggaaagcta
aggactggttttatagacatcactatgaaagtactaatccaaaaataagttcagaagtacacatcccactaggggatgctaaa
ttagtaataacaacatattggggtctgcatacaggagaaagagactggcatttgggtcagggagtctccatagaatggaggaa
aaagagatatagcacacaagtagaccctgacctagcagaccaactaattcatctgcactattttgattgtttttcagaatctg
ctataagaaataccatattaggacgtatagttagtcctaggtgtgaatatcaagcaggacataacaaggtaggatctctacag
tacttggcactagcagcattaataaaaccaaaacagataaagccacctttgcctagtgttaggaaactgacagaggacagatg
gaacaagccccagaagaccaagggccacagagggagccatacaatgaatggacactagagcttttagaggaacttaagagtga
agctgttagacattttcctaggatatggctccataacttaggacaacatatctatgaaacttacggggatacttgggcaggag
tggaagccataataagaattctgcaacaactgctgtttatccatttcagaattgggtgtcgacatagcagaataggcgttact
cgacagaggagagcaagaaatggagccagtagatcctagactagagccctggaagcatccaggaagtcagcctaaaactgctt
gtaccaattgctattgtaaaaagtgttgctttcattgccaagtttgtttcatgacaaaagccttaggcatctcctatggcagg
aagaagcggagacagcgacgaagagctcatcagaacagtcagactcatcaagcttctctatcaaagcagtaagtagtacatgt
aatgcaacctataatagtagcaatagtagcattagtagtagcaataataatagcaatagttgtgtggtccatagtaatcatag
aatataggaaaatattaagacaaagaaaaatagacaggttaattgatagactaatagaaagagcagaagacagtggcaacgag
agtgaaggagaagtatcagcacttgtggagatgggggtggaaatggggcaccacgctccttgggatattgacgatctgtaggc
tagccgtaacgatgttcatcaaatattactgggctgctattaacaagagatggtggtaataacaacaatgggtccgagatctt
cagacctggaggaggcgatatgagggacaattggagaagtgaattatataaatataaagtagtaaaaattgaaccattaggag
tagcacccaccaaggcaaagagaagagtggtgcagagagaaaaaagagcagtgggaataggagctttgttccttgggttcttg
ggagcagcaggaagcactatgggcgcagcgtcaatgacgctgacggtacaggccagacaattattgtctgatatagtgcagca
gcagaacaatttgctgagggctattgaggcgcaacagcatctgttgcaactcacagtctggggcatcaaacagctccaggcaa
gaatcctggctgtggaaagatacctaaaggatcaacagctcctggggatttggggttgctctggaaaactcatttgcaccact
gctgtgccttggaatgctagttggagtaataaatctctggaacagatttggaataacatgacctggatggagtgggacagaga
aattaacaattacacaagcttaatacactccttaattgaagaatcgcaaaaccagcaagaaaagaatgaacaagaattattgg
aattagataaatgggcaagtttgtggaattggtttaacataacaaattggctgtggtatataaaattattcataatgatagta
ggaggcttggtaggtttaagaatagtttttgctgtactttctatagtgaatagagttaggcagggatattcaccattatcgtt
tcagacccacctcccaatcccgaggggacccgacaggcccgaaggaatagaagaagaaggtggagagagagacagagacagat
ccattcgattagtgaacggatccttagcacttatctgggacgatctgcggagcctgtgcctcttcagctaccaccgcttgaga
gacttactcttgattgtaacgaggattgtggaacttctgggacgcagggggtgggaagccctcaaatattggtggaatctcct
acagtattggagtcaggaactaaagaatagtgctgttaacttgctcaatgccacagccatagcagtagctgaggggacagata
gggttatagaagtattacaagcagcttatagagctattcgccacatacctagaagaataagacagggcttggaaaggattttg
ctataagatgggggggaaatggtctaagtcttccgtgataggctggcccgctgttcgagaaaggatgcgccgcgccgaaccag
cggccgatggcgtaggtgccgtatctagggaccttgagaagcacggtgccataacgtcttctaacaccgcagccaacaatgcg
gcatgtgcgtggttggaggcacaggaagaggaggaggtaggattcccggtaacgccacaagtacccttgcgccctatgaccta
caaggccgccgtggacctgagccatttcctgaaggagaaaggtggcttggaagggcttatccactcccaaagacgccaggata
ttcttgacctttggatctaccacacacagggttactttcccgactggcaaaattatacgcccggtcccggtgtacggtatcct
cttacttttgggtggtgctataaactcgtgccggtggagccagataaggtagaggaggccaataaaggtgagaacacaagttt
gcttcaccccgtttcccttcatggtatggacgacccagaacgggaagttcttgaatggcggttcgatagcaggttggcatttc
atcatgtcgcacgggagcttcaccccgagtactttaaaaattgtgcggccgcaggaagcggagctactaacttcagcctgctg
aagcaggctggagacgtggaggagaaccctggacctatgggctggtcatgtatcattctgtttctggtcgcaaccgcaactgg
agtgcattcacaggtgcagctcagcgctgggtcagggatggacgaaaagacaaccggatggcgaggaggacacgtggtcgagg
gactggcaggagagctggaacagctgcgggctagactggaacaccatcctcagggacagcgagagcccggaagtggagcgatc
gcgaaggaagcctgccccacagggctgtacactcattctggagaatgctgtaaagcttgtaacctgggagagggagtggcaca
gccatgcggagccaatcagactgtgtgcgagccttgtctggactccgtcacattctctgatgtggtcagtgccacagaacctt
gcaagccatgtactgagtgcgtgggcctgcagtctatgagtgctccttgtgtggaggctgacgatgcagtctgccggtgtgca
tacggatactatcaggacgagactaccggcagatgtgaagcttgcagggtgtgtgaggcaggctcagggctggtctttagctg
ccaggataaacagaacaccgtgtgcgaggaatgtcctgacgggacatatagcgatgaggccaatcacgtggacccctgcctgc
cttgtactgtgtgcgaggataccgaaaggcagctgcgcgaatgtaccagatgggcagacgccgagtgcgaggaaatcccaggg
cgatggattactcggtccaccccccctgaaggatcagacagcaccgcaccatctacacaggagccagaagcaccaccagagca
ggatctgatcgcctccaccgtggctggcgtggtcacaactgtcatggggagctctcagccagtggtcacccggggcaccacag
ataacctgattcccgtgtattgctccatcctggcagccgtcgtcgtcggactggtggcatacatcgccttcaagcggtgactc
gagacctagaaaaacatggagcaatcacaagtagcaatacagcagctaacaatgctgcttgtgcctggctagaagcacaagag
gaggaagaggtgggttttccagtcacacctcaggtacctttaagaccaatgacttacaaggcagctgtagatcttagccactt
tttaaaagaaaaggggggactggaagggctaattcactcccaaagaagacaagatatccttgatctgtggatctaccacacac
aaggctacttccctgattggcagaactacacaccagggccaggggtcagatatccactgacctttggatggtgctacaagcta
gtaccagttgagccagataaggtagaagaggccaataaaggagagaacaccagcttgttacaccctgtgagcctgcatggaat
ggatgaccctgagagagaagtgttagagtggaggtttgacagccgcctagcatttcatcacgtggcccgagagctgcatccgg
agtacttcaagaactgctgacatcgagcttgctacaagggactttccgctggggactttccagggaggcgtggcctgggcggg
actggggagtggcgagccctcagatgctgcatataagcagctgctttttgcctgtactgggtctctctggttagaccagatct
gagcctgggagctctctggctaactagggaacccactgcttaagcctcaataaagcttgccttgagtgcttcaagtagtgtgt
gcccgtctgttgtgtgactctggtaactagagatccctcagacccttttagtcagtgtggaaaatctctagcacccaggaggt
agaggttgcagtgagccaagatcgcgccactgcattccagcctgggcaagaaaacaagactgtctaaaataataataataagt
taagggtattaaatatatttatacatggaggtcataaaaatatatatatttgggctgggcgcagtggctcacacctgcgcccg
gccctttgggaggccgaggcaggtggatcacctgagtttgggagttccagaccagcctgaccaacatggagaaaccccttctc
tgtgtatttttagtagattttattttatgtgtattttattcacaggtatttctggaaaactgaaactgtttttcctctactct
gataccacaagaatcatcagcacagaggaagacttctgtgatcaaatgtggtgggagagggaggttttcaccagcacatgagc
agtcagttctgccgcagactcggcgggtgtccttcggttcagttccaacaccgcctgcctggagagaggtcagaccacagggt
gagggctcagtccccaagacataaacacccaagacataaacacccaacaggtccaccccgcctgctgcccaggcagagccgat
tcaccaagacgggaattaggatagagaaagagtaagtcacacagagccggctgtgcgggagaacggagttctattatgactca
aatcagtctccccaagcattcggggatcagagtttttaaggataacttagtgtgtagggggccagtgagttggagatgaaagc
gtagggagtcgaaggtgtccttttgcgccgagtcagttcctgggtgggggccacaagatcggatgagccagtttatcaatccg
ggggtgccagctgatccatggagtgcagggtctgcaaaatatctcaagcactgattgatcttaggttttacaatagtgatgtt
accccaggaacaatttggggaaggtcagaatcttgtagcctgtagctgcatgactcctaaaccataatttcttttttgttttt
ttttttttatttttgagacagggtctcactctgtcacctaggctggagtgcagtggtgcaatcacagctcactgcagcctcaa
cgtcgtaagctcaagcgatcctcccacctcagcctgcctggtagctgagactacaagcgacgccccagttaatttttgtattt
ttggtagaggcagcgttttgccgtgtggccctggctggtctcgaactcctgggctcaagtgatccagcctcagcctcccaaag
tgctgggacaaccggggccagtcactgcacctggccctaaaccataatttctaatcttttggctaatttgttagtcctacaaa
ggcagtctagtccccaggcaaaaagggggtttgtttcgggaaagggctgttactgtctttgtttcaaactataaactaagttc
ctcctaaacttagttcggcctacacccaggaatgaacaaggagagcttggaggttagaagcacgatggaattggttaggtcag
atctctttcactgtctgagttataattttgcaatggtggttcaaagactgcccgcttctgacaccagtcgctgcattaatgaa
tcggccaacgcgcggggagaggcggtttgcgtattgggcgctcttccgcttcctcgctcactgactcgctgcgctcggtcgtt
cggctgcggcgagcggtatcagctcactcaaaggcggtaatacggttatccacagaatcaggggataacgcaggaaagaacat
gtgagcaaaaggccagcaaaaggccaggaaccgtaaaaaggccgcgttgctggcgtttttccataggctccgcccccctgacg
agcatcacaaaaatcgacgctcaagtcagaggtggcgaaacccgacaggactataaagataccaggcgtttccccctggaagc
tccctcgtgcgctctcctgttccgaccctgccgcttaccggatacctgtccgcctttctcccttcgggaagcgtggcgctttc
tcatagctcacgctgtaggtatctcagttcggtgtaggtcgttcgctccaagctgggctgtgtgcacgaaccccccgttcagc
ccgaccgctgcgccttatccggtaactatcgtcttgagtccaacccggtaagacacgacttatcgccactggcagcagccact
ggtaacaggattagcagagcgaggtatgtaggcggtgctacagagttcttgaagtggtggcctaactacggctacactagaag
aacagtatttggtatctgcgctctgctgaagccagttaccttcggaaaaagagttggtagctcttgatccggcaaacaaacca
ccgctggtagcggtggtttttttgtttgcaagcagcagattacgcgcagaaaaaaaggatctcaagaagatcctttgatcttt
tctacggggtctgacgctcagtggaacgaaaactcacgttaagggattttggtcatgagattatcaaaaaggatcttcaccta
gatccttttaaattaaaaatgaagttttaaatcaatctaaagtatatatgagtaaacttggtctgacagttaccaatgcttaa
tcagtgaggcacctatctcagcgatctgtctatttcgttcatccatagttgcctgactccccgtcgtgtagataactacgata
cgggagggcttaccatctggccccagtgctgcaatgataccgcgagacccacgctcaccggctccagatttatcagcaataaa
ccagccagccggaagggccgagcgcagaagtggtcctgcaactttatccgcctccatccagtctattaattgttgccgggaag
ctagagtaagtagttcgccagttaatagtttgcgcaacgttgttgccattgctacaggcatcgtggtgtcacgctcgtcgttt
ggtatggcttcattcagctccggttcccaacgatcaaggcgagttacatgatcccccatgttgtgcaaaaaagcggttagctc
cttcggtcctccgatcgttgtcagaagtaagttggccgcagtgttatcactcatggttatggcagcactgcataattctctta
ctgtcatgccatccgtaagatgcttttctgtgactggtgagtactcaaccaagtcattctgagaatagtgtatgcggcgaccg
agttgctcttgcccggcgtcaatacgggataataccgcgccacatagcagaactttaaaagtgctcatcattggaaaacgttc
ttcggggcgaaaactctcaaggatcttaccgctgttgagatccagttcgatgtaacccactcgtgcacccaactgatcttcag
catcttttactttcaccagcgtttctgggtgagcaaaaacaggaaggcaaaatgccgcaaaaaagggaataagggcgacacgg
aaatgttgaatactcatactcttcctttttcaatattattgaagcatttatcagggttattgtctcatgagcggatacatatt
tgaatgtatttagaaaaataaacaaataggggttccgcgcacatttccccgaaaagtgccacctgacgtctaagaaaccatta
ttatcatgacattaacctataaaaataggcgtatcacgaggccctttcgtctcgcgcgtttcggtgatgacggtgaaaacctc
tgacacatgcagctcccggagacggtcacagcttgtctgtaagcggatgccgggagcagacaagcccgtcagggcgcgtcagc
gggtgttggcgggtgtcggggctggcttaactatgcggcatcagagcagattgtactgagagtgcaccatatgcggtgtgaaa
taccgcacagatgcgtaaggagaaaataccgcatcaggcgccattcgccattcaggctgcgcaactgttgggaagggcgatcg
gtgcgggcctcttcgctattacgccaggggaggcagagattgcagtaagctgagatcgcagcactgcactccagcctgggcga
cagagtaagactctgtctcaaaaataaaataaataaatcaatcagatattccaatcttttcctttatttatttatttattttc
tattttggaaacacagtccttccttattccagaattacacatatattctatttttctttatatgctccagttttttttagacc
ttcacctgaaatgtgtgtatacaaaatctaggccagtccagcagagcctaaaggtaaaaaataaaataataaaaaataaataa
aatctagctcactccttcacatcaaaatggagatacagctgttagcattaaataccaaataacccatcttgtcctcaataatt
ttaagcgcctctctccaccacatctaactcctgtcaaaggcatgtgccccttccgggcgctctgctgtgctgccaaccaactg
gcatgtggactctgcagggtccctaactgccaagccccacagtgtgccctgaggctgccccttccttctagcggctgccccca
ctcggctttgctttccctagtttcagttacttgcgttcagccaaggtctgaaactaggtgcgcacagagcggtaagactgcga
gagaaagagaccagctttacagggggtttatcacagtgcaccctgacagtcgtcagcctcacagggggtttatcacattgcac
cctgacagtcgtcagcctcacagggggtttatcacagtgcacccttacaatcattccatttgattcacaatttttttagtctc
tactgtgcctaacttgtaagttaaatttgatcagaggtgtgttcccagaggggaaaacagtatatacagggttcagtactatc
gcatttcaggcctccacctgggtcttggaatgtgtcccccgaggggtgatgactacctcagttggatctccacaggtcacagt
gacacaagataaccaagacacctcccaaggctaccacaatgggccgccctccacgtgcacatggccggaggaactgccatgtc
ggaggtgcaagcacacctgcgcatcagagtccttggtgtggagggagggaccagcgcagcttccagccatccacctgatgaac
agaacctagggaaagccccagttctacttacaccaggaaaggc
~~~

